# Arsenic-Driven Trafficking of ACR3 Transporters Is Conserved across Plant Lineages

**DOI:** 10.64898/2025.12.14.694196

**Authors:** Kacper Zbieralski, Alicja Dolzblasz, Paulina Tomaszewska, Katarzyna Mizio, Carmen Mata, Verena Kriechbaumer, Anna Janiczek, Jacek Staszewski, Ignacy Bonter, Robert Wysocki, Ewa Maciaszczyk-Dziubinska, Donata Wawrzycka

## Abstract

Arsenic contamination poses a major environmental and public health challenge. In liverworts and ferns, ACR3 transporters mediate arsenite efflux and contribute to arsenic tolerance. We recently showed that ACR3 from *Marchantia polymorpha* (MpACR3) contains an extended N-terminal region that couples arsenic sensing to membrane trafficking. However, the function and regulation of ACR3 transporters across photosynthetic lineages remain unclear. Here, we characterize ACR3 transporters from phylogenetically distant algae and land plants using heterologous expression in yeast. All examined ACR3 proteins function as arsenic efflux transporters but display two distinct trafficking modes: constitutive plasma membrane localization or metalloid-driven relocalization from the endomembrane system. Like the previously characterized MpACR3, we show that this inducible trafficking is governed by an extended N-terminal domain unique to plant ACR3 proteins, integrating a conserved di-arginine ER/Golgi retention signal with cysteine-based arsenic sensing. These findings establish a direct mechanistic link between metalloid perception and subcellular targeting. Functional assays in *Arabidopsis thaliana* demonstrated that *Physcomitrium patens* ACR3 confers arsenic tolerance, reduces root arsenic accumulation, and retains metalloid-dependent trafficking, supporting its physiological relevance. Together, our findings uncover a conserved mechanism linking arsenic sensing to ACR3 trafficking and provide new insight into the evolution of arsenic tolerance in plants.

**Summary statement:** Plant ACR3 transporters mediate arsenic efflux and exhibit distinct trafficking modes. An N-terminal domain links arsenic sensing to the endomembrane system-plasma membrane relocalization, revealing a conserved mechanism controlling ACR3 function and arsenic tolerance across plant lineages.

## 1. Introduction

Contamination of the environment with arsenic, a highly toxic metalloid, has emerged as a significant global health and environmental concern. Chronic exposure to prevalent inorganic arsenic forms, trivalent arsenite [As(OH)₃, As(III)] and pentavalent arsenate [H₂AsO₄⁻/HAsO₄²⁻, As(V)], has been repeatedly associated with severe health effects, including cancer, neurodegenerative disorders, abnormalities in cognitive development, and cardiovascular diseases. Arsenic is naturally distributed throughout the earth’s crust and can leach into water supplies, while anthropogenic activities such as mining, agriculture, and industrial processes further amplify the risk of contamination (Ganie et al. 2024). Current estimates suggest that up to 200 million people worldwide may be chronically exposed to unsafe arsenic levels (Podgorski and Berg 2020). The main routes of arsenic consumption by humans are contaminated groundwater and food, particularly agricultural produce (Podgorski and Berg 2020; Mawia et al. 2021; Molin et al. 2015). Thus, reducing arsenic contamination of food, water reservoirs and farmlands is of utmost importance from both ecological and human health perspectives.

A growing body of evidence highlights the pivotal role of the ACR3 family of arsenic transporters in conferring arsenic tolerance in plants. The transporters have been mostly characterized in budding yeast and bacteria, where they act as As(III) exporters at the plasma membrane (PM) (Bobrowicz et al. 1997; Wysocki et al. 1997; Sato and Kobayashi 1998; Ghosh et al. 1999; Fu et al. 2009; Maciaszczyk-Dziubinska et al. 2010; Villadangos et al. 2012). In plants, however, the biological functions of ACR3 orthologs remain insufficiently described.

The ACR3 family is a member of the bile/arsenite/riboflavin transporter (BART) superfamily and a distant relative to the bile acid sodium symporters (BASS/SBF family). These families share a 10-transmembrane (TM) topology with a Na^+^/H^+^ antiporter-like fold but have diverged substantially in substrate specificity and transport mechanisms (Mansour et al. 2007). ACR3 proteins are As(III)/H^+^ antiporters, whose activity largely depends on highly conserved cysteine and glutamate residues (Villadangos et al. 2012; Maciaszczyk-Dziubinska et al. 2011; Markowska et al. 2015; Wawrzycka et al. 2017). Although plant orthologs are predicted to share a conserved ACR3 topology (Indriolo et al. 2010; Li et al. 2024; Mizio et al. 2025), the functionality of most plant family members, as well as the corresponding residues required for metalloid transport, has not yet been experimentally confirmed.

In plants, ACR3 transporters were first characterized in the arsenic hyperaccumulator *Pteris vittata*, which encodes five paralogs (PvACR3, PvACR3;1, PvACR3;2, PvACR3;3, and PvACR3;4) (Indriolo et al. 2010; Chen et al. 2013; Chen et al. 2017; Chen et al. 2021; Wang et al. 2018; Sun et al. 2023). This enables the fern to accumulate high arsenic levels, with concentrations ranging from 66-6,151 mg/kg in soil-based experiments to 22,630-27,000 mg/kg under hydroponic conditions (Wang et al. 2002; Gonzaga et al. 2008). Moreover, *PvACR3*-RNAi gametophytes of *P. vittata* exhibit arsenic-sensitive growth (Indriolo et al. 2010). Other *Pteris* species also express multiple *ACR3* transporters, including PcACR3 and PcACR3;1 in *P. cretica* and PmACR3 and PmACR3;1 in *P. multifida* (Popov et al. 2021; Li et al. 2025), while the non-hyperaccumulator *P. ensiformis* expresses at least one ACR3 protein (PeACR3) (Sun et al. 2023).

Recent functional studies in the model bryophyte *Marchantia polymorpha* revealed that its native ACR3 (MpACR3) enhances arsenic tolerance and reduces accumulation in plant tissues when overexpressed, whereas loss-of-function mutants are hypersensitive to arsenic (Li et al. 2024; Mizio et al. 2025). Cross-species complementation assays further confirm conservation of ACR3 function, as MpACR3, PvACR3, PvACR3;1, PvACR3;2, PvACR3;3, and PcACR3 at least partially complement the hypersensitivity phenotype of budding yeast cells lacking the endogenous *ACR3* gene (*ScACR3*) (Indriolo et al. 2010; Chen et al. 2013; Chen et al. 2017; Chen et al. 2021; Popov et al. 2021; Dutta et al. 2024). Similarly, when heterologously expressed in the *ACR3*-lacking model alga *Chlamydomonas reinhardtii*, PvACR3 enhanced arsenic export in cells exposed to arsenate in low-phosphate medium (Ramírez-Rodríguez et al. 2019). Notably, the acidophilic alga *Chlamydomonas eustigma*, which encodes a putative ACR3 transporter (CeACR3), tolerates arsenate levels over tenfold higher than *C. reinhardtii*, indicating the functionality of CeACR3 (Hirooka et al. 2017).

Although missing in flowering plants, several heterologously expressed ACR3 transporters effectively improved arsenic tolerance in transgenic *Arabidopsis thaliana*, *Nicotiana tabacum* and *Oryza sativa* lines. For example, expression of ScACR3, PvACR3, and PvACR3;2 reduced overall arsenic accumulation while enhancing root-to-shoot arsenic translocation in *A. thaliana* and tobacco. (Ali et al. 2012; Chen et al. 2013; Chen et al. 2021). On the other hand, *PvACR3;1* expression decreased arsenic translocation factor and increased arsenic accumulation in roots of these model plants (Chen et al. 2017), and *PvACR3;3* increased arsenic root accumulation in tobacco (Chen et al. 2021). In *O. sativa*, PvACR3;1 also reduced arsenic accumulation in rice grains (Chen et al. 2019), while ScACR3 lowered arsenic accumulation in straws, grains, and roots without altering its root-to-shoot transport (Duan et al. 2012). Although the effect of MpACR3 heterologous expression on arsenic allocation in angiosperm tissues has not been determined so far, MpACR3 significantly decreased total arsenic accumulation in *A*. *thaliana* seedlings (Mizio et al. 2025).

Current evidence highlights functional and regulatory divergence of ACR3 proteins across species. In yeast and bacteria, *ACR3* expression is generally low at baseline but activated during arsenic stress by induction in yeast or de-repression in bacteria (Wysocki et al. 2004; Maciaszczyk-Dziubinska et al. 2010; Branco et al., 2008; Rawle et al. 2021). In plants, *ACR3* expression patterns are more diverse, varying with tissue type, species, and arsenic exposure. For example, *MpACR3* in *M. polymorpha* is constitutively expressed with moderate induction by arsenic (Li et al. 2024; Dutta et al. 2024; Mizio et al. 2025), while *C. eustigma* maintains relatively high basal *CeACR3* expression (Hirooka et al. 2017). In *P. vittata*, *PvACR3* is strongly arsenic-inducible in roots and gametophytes, whereas *PvACR3;1* is constitutively expressed and arsenic-insensitive (Indriolo et al. 2010). Several additional *PvACR3* paralogs show primarily frond-located expression, with *PvACR3;2* being arsenic-insensitive and *PvACR3;3/4* arsenic-inducible (Chen et al. 2021; Sun et al. 2023). Other *Pteris* species (e.g., *P. cretica* and *P. ensiformis*) also display arsenic-inducible ACR3 variants (Popov et al. 2021; Sun et al. 2023).

Beyond transcriptional control, plant ACR3s exhibit distinct subcellular localizations consistent with divergent functions. In *P. vittata*, PvACR3 and PvACR3;3 localize to the vacuolar membrane (VM) to mediate arsenic sequestration into the vacuole (Indriolo et al. 2010; Chen et al. 2017; Chen et al. 2019; Chen et al. 2021; Sun et al. 2023), whereas PvACR3;2 functions as a PM transporter for root-to-shoot arsenic translocation (Chen et al. 2021; Sun et al. 2023). When heterologously expressed in *A. thaliana* and rice, PvACR3;1 localized to the VM and facilitated increased arsenic accumulation in roots (Chen et al. 2017; Chen et al. 2019), while recent studies suggest that PvACR3;4 may also localize to the VM in *P. vittata* frond cells (Sun et al. 2023). Heterologous expression studies corroborate host-dependent protein trafficking, with PvACR3 localizing to the PM and mediating arsenic efflux in yeast and *A. thaliana* (Chen et al. 2017; Wang et al. 2018; Chen et al. 2021) and PvACR3;1 supporting efflux activity in yeast, suggesting possible plasmalemmal localization (Chen et al. 2017).

Unlike previously characterized ACR3 orthologs, MpACR3 localizes to both the PM and intracellular compartments. Specifically, it is found in the Golgi network in *M. polymorpha* and *A. thaliana*, and in the endoplasmic reticulum (ER) in yeast. Moreover, MpACR3 undergoes arsenic-dependent redistribution from the Golgi/ER to the PM (Mizio et al. 2025). This process is mediated by an N-terminal cytosolic metalloid-sensing domain which is predicted to coordinate arsenic via three conserved cysteine residues, one of which resides within an arginine-based R-C-R motif required for intracellular retention (Mizio et al. 2025). We previously proposed that arsenic relieves intracellular retention of MpACR3, likely through conformational changes in the N-terminal domain that alter protein-protein interactions and/or post-translational modifications, ultimately leading to its subcellular redistribution (Mizio et al. 2025).

Notably, plant ACR3 orthologs with extended N-terminal regions are predicted to harbor similar metalloid-sensing domains containing a highly conserved Φ_1_-R-C-R-Φ_2_ motif (where Φ denotes a bulky hydrophobic residue), hereafter referred to as the di-arginine motif. Within this motif, the two arginine residues and the Φ_1_ residue contribute to intracellular retention, whereas the intervening cysteine and the Φ_2_ residue are required for arsenic-dependent release from the endomembrane system (Mizio et al. 2025). However, it remains unclear whether di-arginine motif-dependent intracellular retention and arsenic-responsive trafficking are conserved and functionally relevant across other plant ACR3 orthologs.

To date, little information on plant ACR3 orthologs outside of *Pteris* genus and *M. polymorpha* species is available; thus, our understanding of plant ACR3 evolution and diversity remains elusive. To test whether functional conservation in plant ACR3 transporters extends across distinct environmental challenges, we used *S. cerevisiae* as a heterologous expression system to characterize previously unstudied plant ACR3 transporters. Based on our updated phylogenetic analysis, we selected unicellular green algae, including psychrophilic *Coccomyxa subellipsoidea* (CsACR3), a common bioindicator *Raphidocelis subcapitata* (RsACR3), and acidophilic, heavy metal-tolerant *Chlamydomonas eustigma* (CeACR3), a model moss *Physcomitrium patens* (PpACR3), and an emerging gymnosperm model *Picea sitchensis* (PsACR3). The proteins were characterized alongside the previously described ACR3 proteins from the model liverwort *M. polymorpha* (MpACR3) and the arsenic-hyperaccumulating fern *P. vittata* (PvACR3).

First, we show that the selected proteins are predicted to share the common ACR3 topology, with CeACR3, MpACR3, PpACR3 and PsACR3 equipped with exceptionally elongated N-termini of unknown function. Second, we demonstrate that the selected genes encode functional arsenic transporters that confer differential resistance to arsenic compounds in yeast. Third, we show that the arsenic-dependent PpACR3 trafficking regulation is functional in *A*. *thaliana*, and the transporter enhances growth under arsenic stress by reducing arsenic accumulation. Finally, we prove that N-terminal domains in CeACR3, PpACR3, and PsACR3 regulate the conserved mechanism of their intracellular retention and arsenic-dependent relocalization to the PM. These findings provide new insight into the evolution and functional characteristics of previously undescribed plant ACR3 transporters, expanding on current knowledge of cellular arsenic detoxification mechanisms in plants.

## 2. Results

### 2.1. Conservation of ACR3 Transporters across Distant Plant Lineages

Phylogenetic analysis of plant ACR3 transporters revealed two distinct clades corresponding to algal and land plant ACR3 proteins, consistent with the current understanding of plant evolutionary history (Figure 1). Strikingly, an exception is the ACR3 ortholog from the lycophyte *Selaginella moellendorffii* (SmACR3), which is positioned at the base of the land plant clade. Moreover, ACR3 orthologs are widely distributed across chlorophytes, bryophytes, lycophytes, ferns, and gymnosperms (Figure 1; Table S1). In contrast, ACR3 orthologs are absent from the prasinodermophyte *Prasinoderma coloniale*, several chlorophyte green algae, including the model species *Chlamydomonas reinhardtii* and *Volvox carteri*, as well as the charophyte green algae *Chara braunii*, *Chlorokybus atmophyticus*, and *Mesostigma viride*.

**Figure 1.**
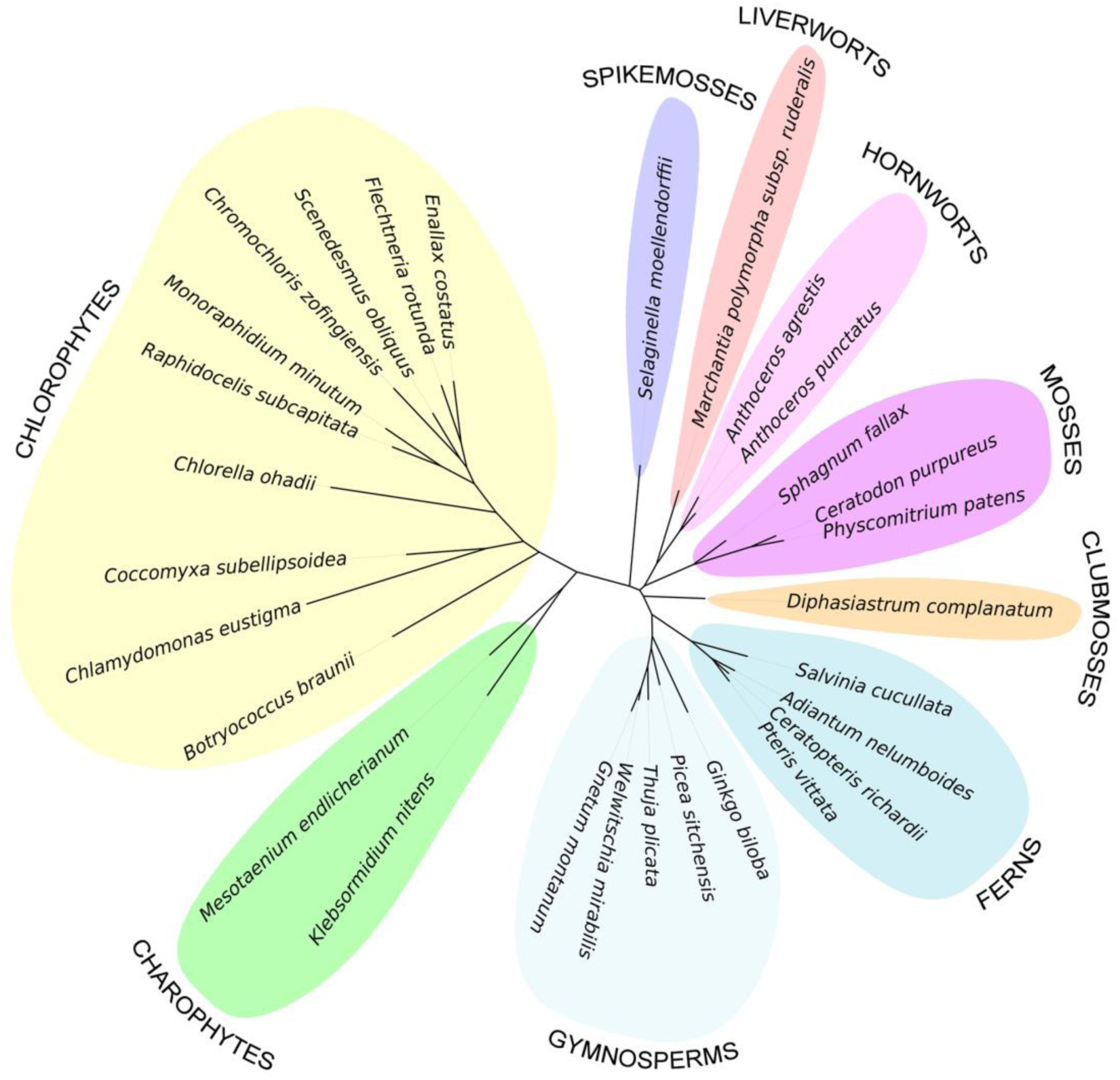
Phylogenetic tree of the plant members of the ACR3 family. The sequences were aligned using ClustalW 2.1. The phylogenetic relationship was inferred by the maximum likelihood method. 1000 bootstrap replicates were generated to estimate branch support values. Sequence details are provided in Table S1.

### 2.2. Unique N-terminal Extensions Are a Prevalent Feature of Plant ACR3 Orthologs

So far, plant ACR3 transporters have only been functionally characterized in *Pteris* ferns and the liverwort *M. polymorpha* (Indriolo et al. 2010; Chen et al. 2013; Chen et al. 2017; Wang et al., 2018; Chen et al., 2021; Sun et al. 2023; Li et al. 2024, Dutta et al. 2024; Mizio et al. 2025). Consequently, current understanding of the evolutionary diversity of plant ACR3s remains restricted to these taxa, despite the family prevalence in plant kingdom (Figure 1; Table S1). In order to elucidate the properties of previously uncharacterized plant ACR3 transporters, we selected a set of *ACR3* genes from evolutionarily and ecologically divergent species: green algae *C. subellipsoidea* C-169 (*CsACR3*), *R. subcapitata* (*RsACR3*), *C. eustigma* (*CeACR3*), moss *P. patens* (*PpACR3*), and conifer *P. sitchensis* (*PsACR3*) for functional analysis alongside previously described genes from liverwort *M. polymorpha* (*MpACR3*) (Dutta et al. 2024, Li et al. 2024; Mizio et al. 2025) and fern *P. vittata* (*PvACR3*) (Indriolo et al. 2010) (Tables S1, S2).

The *ACR3* genes encode proteins of varying lengths, ranging from 360 aa (CsACR3) to 495 aa (CeACR3), with a mean sequence identity of 54.3% and relatively high mean similarity of 70.3% (Table S3). Multiple sequence alignment followed by topology predictions indicate that the putative transporters are membrane proteins sharing the ACR3 topology of 10 transmembrane (TM) regions, with both termini oriented towards the cytoplasm (Figures S1, S2; Table S4). Importantly, all proteins retain the functionally critical cysteine residue in TM4 and the glutamate residues in TM9-10, previously shown to be essential for arsenite transport in yeast and bacterial ACR3s (Villadangos et al. 2012; Maciaszczyk-Dziubinska et al. 2011; Markowska et al. 2015; Wawrzycka et al. 2017) (Figure S1). Notably, while the transporters showed no significant differences in intra- and extracellular loop lengths, they exhibited either short (CsACR3, RsACR3, and PvACR3) or elongated (CeACR3, MpACR3, PpACR3, and PsACR3), poorly conserved N-terminal tails, with an additional C-terminal extension in CeACR3 (Figures S1, S3). Interestingly, such N-terminal extensions (104- 163 residues) are a common characteristic of plant ACR3 orthologues (Figure S3). Only a few ACR3 sequences, primarily from ferns and certain algae, contain N-terminal tails shorter than 100 residues (Figure S3). Collectively, these results suggest that plant ACR3 orthologs from phylogenetically distant species might perform similar biological functions. However, sufficient sequence variation exists, especially in the N-terminal cytoplasmic regions, to indicate possible adaptations to specific organismal, cellular, or environmental contexts.

### 2.3. Plant *ACR3* Genes Confer Arsenic Tolerance in Yeast

*In silico* analyses strongly suggest that the putative ACR3 orthologs might be functional arsenic transporters (Figures 1, S1, S2). For further analysis, we used the yeast *acr3*Δ mutant lacking the endogenous *ACR3* (*ScACR3*) gene as a heterologous expression system. Consistent with previously generated plasmids pScACR3 (Maciaszczyk-Dziubinska et al. 2014) and pMpACR3 (Mizio et al. 2025), synthetic *CsACR3*, *RsACR3*, *CeACR3*, *PpACR3*, and *PsACR3* genes were cloned and expressed under the control of the constitutive *MET17* promoter as C-terminal GFP fusion constructs, yielding pCsACR3, pRsACR3, pCeACR3, pPpACR3, and pPsACR3, respectively.

All obtained plasmids provided high expression of the analyzed fusion proteins as confirmed by Western blotting (Figure 2A). To test the functionality of the constructs, we performed growth assays on solid media in the presence of various concentrations of As(III) and As(V) using empty vector as a negative control and the pScACR3 construct as a positive control. While all the *ACR3*-bearing plasmids improved the growth of the *acr3*Δ mutant in the presence of As(III), only pMpACR3 and pCsACR3 provided full complementation of the *acr3*Δ phenotype (Figure 2B). In *acr3*Δ cells, As(III) tolerance was markedly enhanced by CeACR3 and PvACR3, moderately by RsACR3 and PpACR3, and only slightly by PsACR3 (Figure 2B). In contrast, the expression of all plant *ACR3s* restored As(V) tolerance in *acr3*Δ cells, with four (RsACR3, PpACR3, PvACR3, PsACR3) slightly outperforming ScAcr3 (Figure 2B). In arsenic-accumulation assays, most ACR3-expressing strains contained less intracellular arsenic versus the empty-vector control (p < 0.05) (Figure 2C, D). RsACR3 and PsACR3, however, performed poorly at high As(III) (1 mM for 2 h), but markedly reduced As(III) accumulation at the lower exposure (0.1 mM for 4 h). Together, these results show that, when expressed in yeast, the analyzed plant ACR3 proteins facilitate the efflux of As(III), albeit with varying efficiencies.

**Figure 2.**
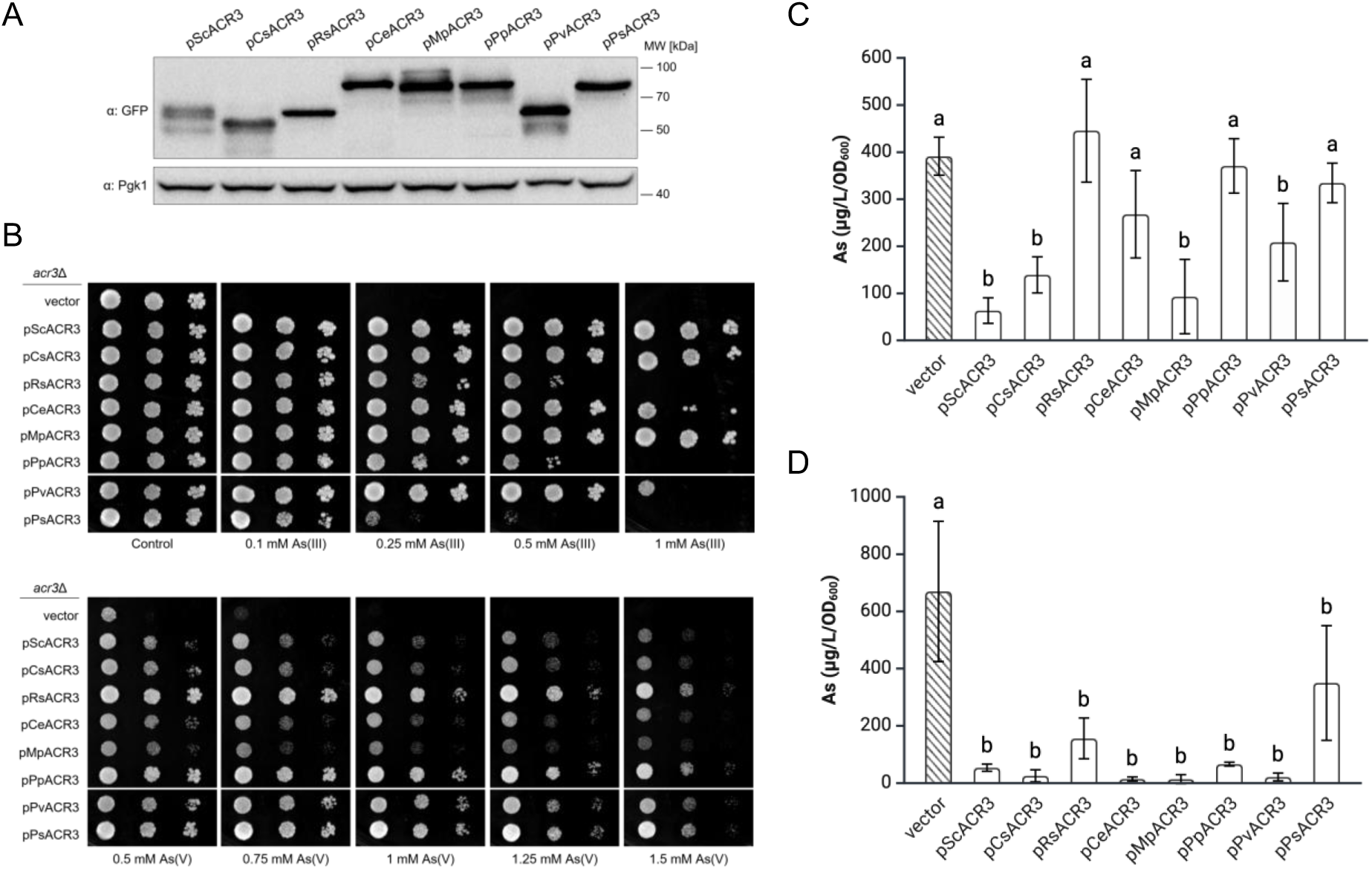
Functional analysis of plant ACR3 proteins in *S. cerevisiae*. (**A**) Western blot analysis of GFP-fused proteins expressed in *acr3*Δ cells. Pgk1 (3-phosphoglycerate kinase) was used as a loading control. MW, molecular weight. (**B**) Growth assays demonstrating that the expression of selected plant *ACR3* genes rescues the sensitivity of *acr3*Δ cells to As(III) and As(V). (**C, D**) Metalloid accumulation assays demonstrating arsenic extrusion capability of selected ACR3 orthologs expressed in *acr3*Δ mutant after exposure to 1 mM As(III) for 2 h (C) or 0.1 mM As(III) for 4 h (**D**). Error bars represent mean values ± standard deviation (SD) from three independent experiments (n = 3) with three technical repeats. Control groups are indicated by diagonal lines. Different letters indicate statistically significant differences determined by one-way ANOVA (p < 0.05).

### 2.4. Arsenic Stress Drives ER-to-PM Relocalization of N-Terminally Extended Plant ACR3 Orthologs

Considering their apparent efflux activity, we hypothesized that the plant ACR3 transporters localize to the yeast PM. To address this, we observed the subcellular localization of the ACR3-GFP fusion proteins in the *acr3*Δ strain using fluorescence microscopy. Under basal conditions, all fusion proteins were detected in intracellular ring-like structures, as well as continuous (CsACR3, RsACR3, and PvACR3) or punctate (CeACR3, MpACR3, PpACR3, and PsACR3) peripheral compartments (Figure 3A). While the uniform signal at the cell periphery corresponds to PM localization, the ragged peripheral and continuous intracellular signals were consistent with cortical and perinuclear ER retention, respectively (Zhu et al. 2019). Co-localization with the ER marker mCherry-HDEL confirmed that under basal conditions CsACR3-GFP, RsACR3-GFP, and PvACR3-GFP reside at both the PM and ER, whereas CeACR3-GFP, MpACR3-GFP, PpACR3-GFP, and PsACR3-GFP are primarily ER-localized (Figure 3A). Nevertheless, other intracellular compartments such as the Golgi apparatus cannot be excluded.

**Figure 3.**
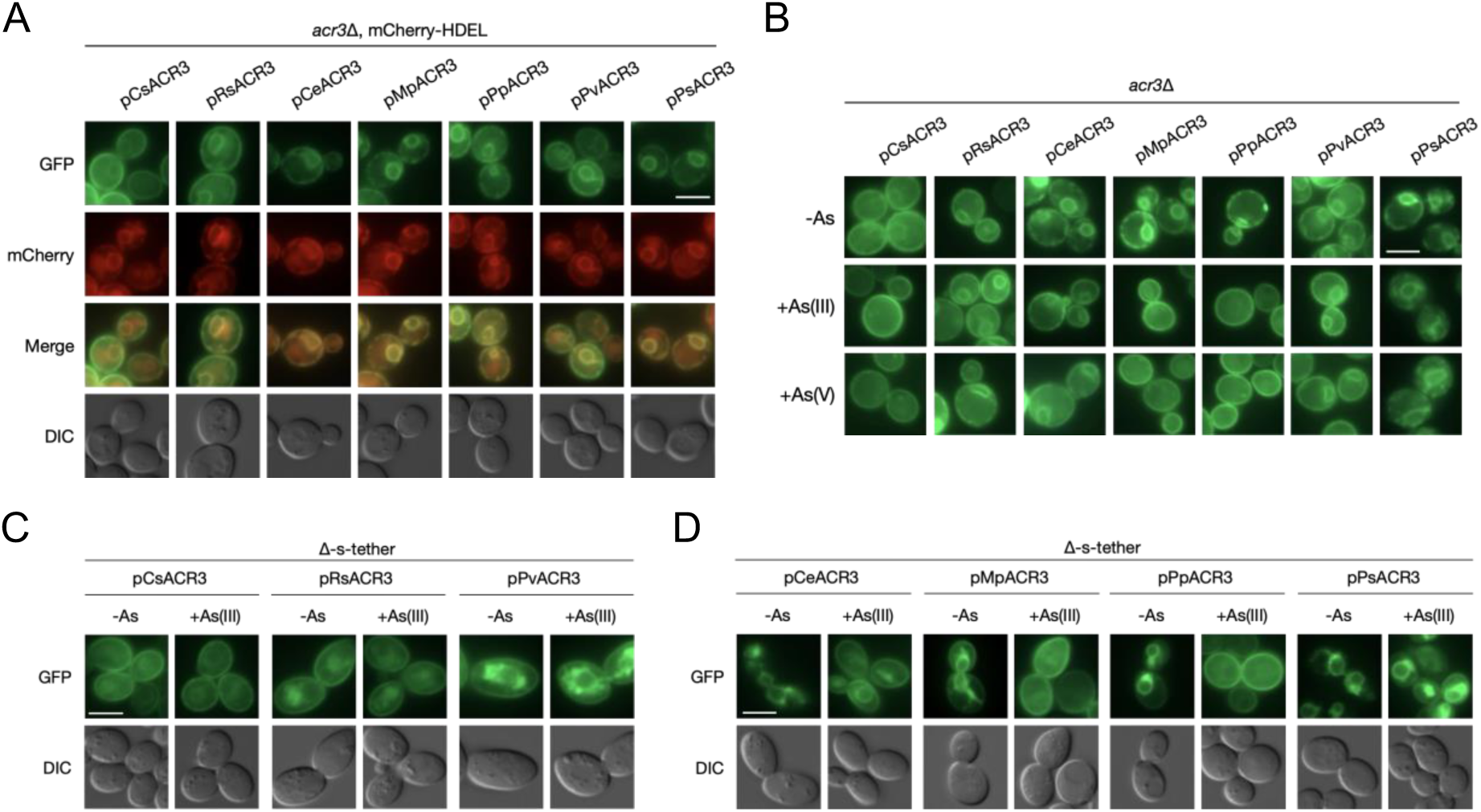
Subcellular localization of plant ACR3 orthologs in *S. cerevisiae*. (**A**) Co-localization of GFP-tagged plant ACR3 proteins with endoplasmic reticulum-localized mCherry in *acr3*Δ cells. (**B**) Localization of selected ACR3 proteins in *acr3*Δ cells under control conditions or after 2 h exposure to 0.1 mM As(III). (**C, D**) Localization of short-tailed (**C**) and long-tailed (**D**) ACR3 orthologs in Δ-s-tether cells under control conditions or following treatment with 0.3 mM As(III) for 2 h, as determined by fluorescent microscopy. Scale bars = 5 µm.

The dominant reticular localization of N-terminally extended ACR3 orthologs did not correspond to their ability to reduce arsenic concentration in yeast cells (Figure 2C, D). Thus, we hypothesized that arsenic exposure might affect the subcellular localization of the plant ACR3-GFP fusions. Indeed, upon exposure to As(III) or As(V), MpACR3-GFP and PpACR3-GFP underwent a pronounced ER-to-PM relocalization, evident as near-complete loss of ER signal with concomitant enrichment at the cell periphery, whereas CsACR3-GFP, RsACR3-GFP, and PvACR3-GFP did not (Figure 3B). Under the same conditions, CeACR3-GFP and PsACR3-GFP retained strong reticular fluorescence, preventing unambiguous confirmation of an ER-to-PM shift (Figure 3B). To clarify this, we further analyzed the localization of the fusion proteins in the mutant strain with the ER separated from the PM due to the lack of all ER-to-PM tethering proteins (Δ-s-tether strain) (Quon et al. 2018). Indeed, we confirmed that CsACR3-GFP, RsACR3-GFP and PvACR3-GFP localized to the ER and the PM regardless of arsenic exposure (Figure 3C). The localization of MpACR3 and PpACR3, on the other hand, changed from primarily reticular to PM upon arsenic treatment (Figure 3D). Nevertheless, a fraction of the GFP signal for these proteins was also observed in the PM even in the absence of the metalloid (Figure 3D). Conversely, under normal conditions, the GFP signal of CeACR3-GFP and PsACR3-GFP was confined to intracellular compartments, whereas arsenic induced strong (CeACR3-GFP) and intermediate (PsACR3-GFP) PM signals (Figure 3D). Together, these results demonstrate that plant ACR3 transporters localize to the PM when expressed in yeast, with the PM targeting of the N-terminally elongated orthologs being at least partially regulated by arsenic exposure.

### 2.5. PpACR3 Expression Enhances Arsenic Resistance in *A. thaliana*

For functional analysis in plants, including arsenic tolerance assays and localization studies, we selected the long-tailed PpACR3 from the well-established bryophyte model *P. patens*, as it exhibited differential resistance to As(III) and As(V) (Figure 2), as well as arsenic-induced relocalization from the ER to the PM in yeast cells (Figure 4).

**Figure 4.**
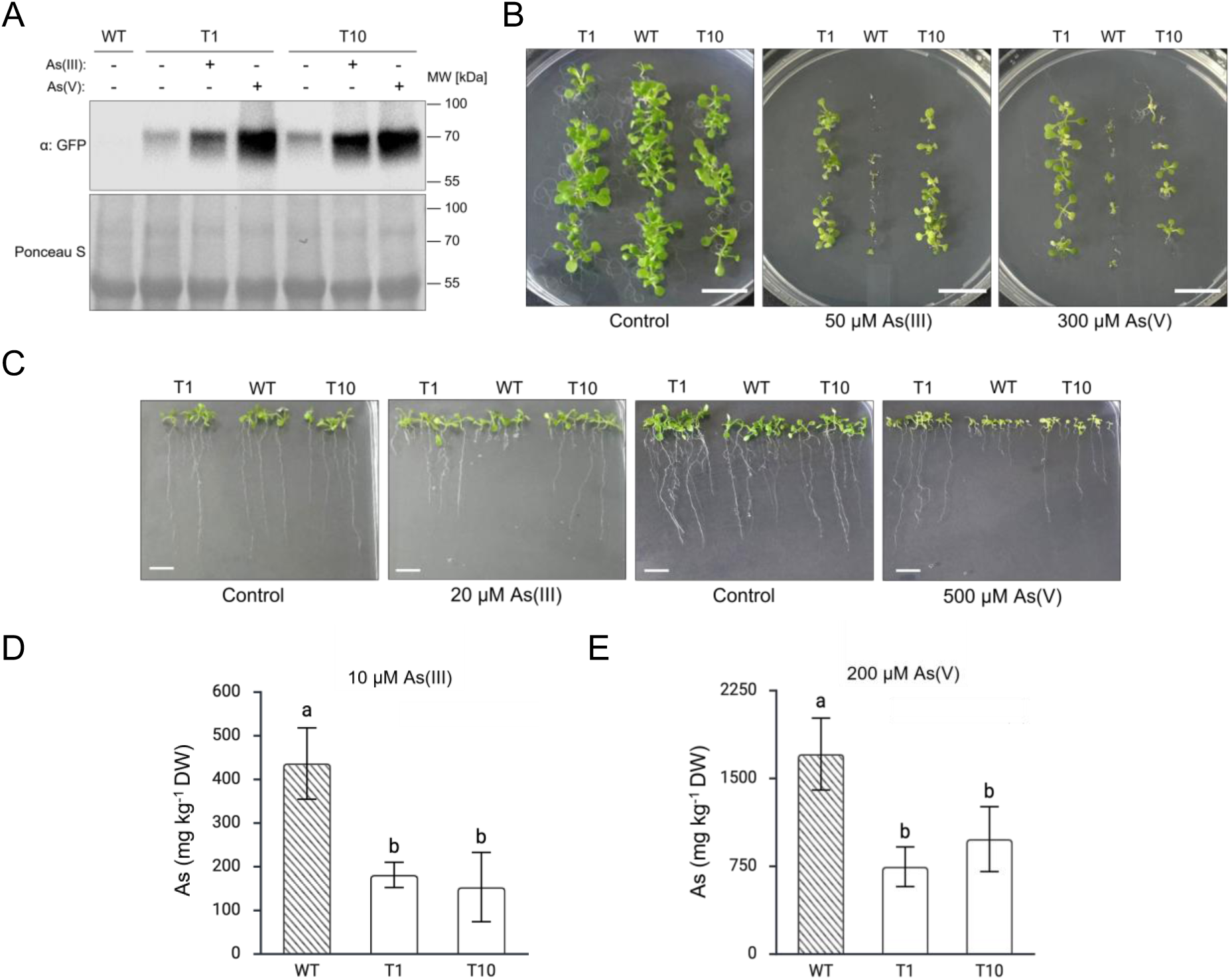
Overexpression of PpACR3 in *A. thaliana*. (**A**) Western blot analysis demonstrates increased accumulation of PpACR3-GFP following treatment with 20 µM As(III) or 500 µM As(V) in two independent pro35S:PpACR3-GFP transgenic lines (T1 and T10). Ponceau S staining was used as a loading control. MW, molecular weight; WT, wild-type. (**B**) Germination of WT and transgenic (T1 and T10 lines) seeds in the absence or presence of 50 µM As(III) or 300 µM As(V). Scale bars = 1 cm. ( **C**) Comparative growth of WT and transgenic (T1 and T10) seedlings grown for 4 days in MS medium followed by 7 days of growth in the absence or presence of As(III) or As(V). Scale bars = 1 cm. (**D, E**) Arsenic accumulation in WT, T1, and T10 seedlings grown for 2 weeks in the presence of As(III) (**D**) or As(V) (**E**). Error bars represent mean values ± standard deviation (SD) from three independent experiments (n = 3). Within each group and treatment condition, 5-10 seedlings were pooled for arsenic content determination. Control groups are indicated by diagonal lines. Different letters indicate statistically significant differences determined by one-way ANOVA (p < 0.05). DW, dry weight.

First, we examined whether *PpACR3* expression is influenced by metalloid exposure in wild-type *P. patens* ssp. *patens* (Gransden 2004 ecotype). Using qPCR, we measured *PpACR3* transcript levels in gametophyte tissues of plants grown in the absence or presence of As(III) or As(V) at 100 µM and 200 µM concentrations. *PpACR3* showed constant low expression even in the absence of arsenic (Figure S4). Moreover, *PpACR3* expression was unaffected by exposure to the metalloid in both arsenic species- and dose-dependent manner, indicating arsenic-independent *PpACR3* expression regulation in the native organism (Figure S4).

Next, we transformed wild-type *A. thaliana* Columbia-0 (Col-0) with *PpACR3* fused to *GFP* under the constitutive CaMV 35S promoter. Two independently selected transgenic lines bearing *pro35S:PpACR3-GFP* (T1 and T10) were used for subsequent analysis. No phenotypical differences were observed for soil-grown wild-type, T1 and T10 lines under normal growth conditions (Figure S5). Western blot analysis confirmed PpACR3-GFP expression in both transgenic lines, with markedly higher protein levels detected in plants grown for two weeks on solid medium supplemented with 20 µM As(III) or 300 µM As(V) compared with untreated controls (Figure 4A). Because PpACR3-GFP expression was driven by the constitutive CaMV 35S promoter, the arsenic-dependent increase in protein abundance is unlikely to result from transcriptional regulation and may instead reflect enhanced protein stability and/or accumulation under arsenic stress.

To evaluate the effect of *PpACR3* expression on *A. thaliana* tolerance to arsenic stress, we first analyzed seed germination on As(III)- and As(V)-supplemented solid media. No differences between wild-type and transgenic lines were observed under normal conditions. However, the exposure to As(III) or As(V) significantly inhibited germination of wild-type plants, while both T1 and T10 lines germinated successfully (Figure 4B). Moreover, wild-type seedlings exhibited more severe growth inhibition than T1 and T10 lines when germinated on standard MS medium, transferred to As(III)- and As(V)-containing plates and grown for two weeks (Figure 4C). Consistently, PpACR3 expression reduced arsenic accumulation in plants grown for two weeks on plates containing As(III) or As(V) at concentrations tolerated by wild-type *A. thaliana* (p < 0.05) (Figure 4D). Thus, unlike in yeast cells, PpACR3 conferred similarly high tolerance to both As(III) and As(V) when expressed in *A. thaliana*.

### 2.6. Arsenic Regulates ER-PM Trafficking of PpACR3 in *A. thaliana*

Previously, we demonstrated that MpACR3 trafficking to the plasma membrane (PM) is delayed in the absence of arsenic in both yeast and plant cells. However, MpACR3-GFP exhibits dual localization to the ER and PM in yeast, whereas in plants it localizes predominantly to the Golgi and PM (Mizio et al. 2025). Here, we show that other plant ACR3 proteins with extended N-terminal regions, including PpACR3, also display dual ER/PM localization in yeast cells and accumulate at the PM in response to arsenic stress (Figure 3).

To determine the subcellular localization pattern of PpACR3 in plants, we observed PpACR3-GFP in *A. thaliana* root cells using Lattice Lightsheet LL7 microscope. Under normal conditions, strong intracellular and peripheral GFP signals were detected (Figures 5A, S6A). However, exposure to 20 µM As(III) or 500 µM As(V) enhanced peripheral and diminished intracellular signals, suggesting arsenic-dependent regulation of PpACR3 trafficking (Figures 5B, C, S6B, C). Cross sections of individual cells suggested ER localization of PpACR3-GFP under normal conditions, as evidenced by the characteristic perinuclear signal observed in some root cells (Figures 5A, S6A). The signal was no longer detectable following arsenic exposure, further supporting arsenic-induced relocalization of PpACR3 from the endomembrane system to the PM (Figure 5B, C).

**Figure 5.**
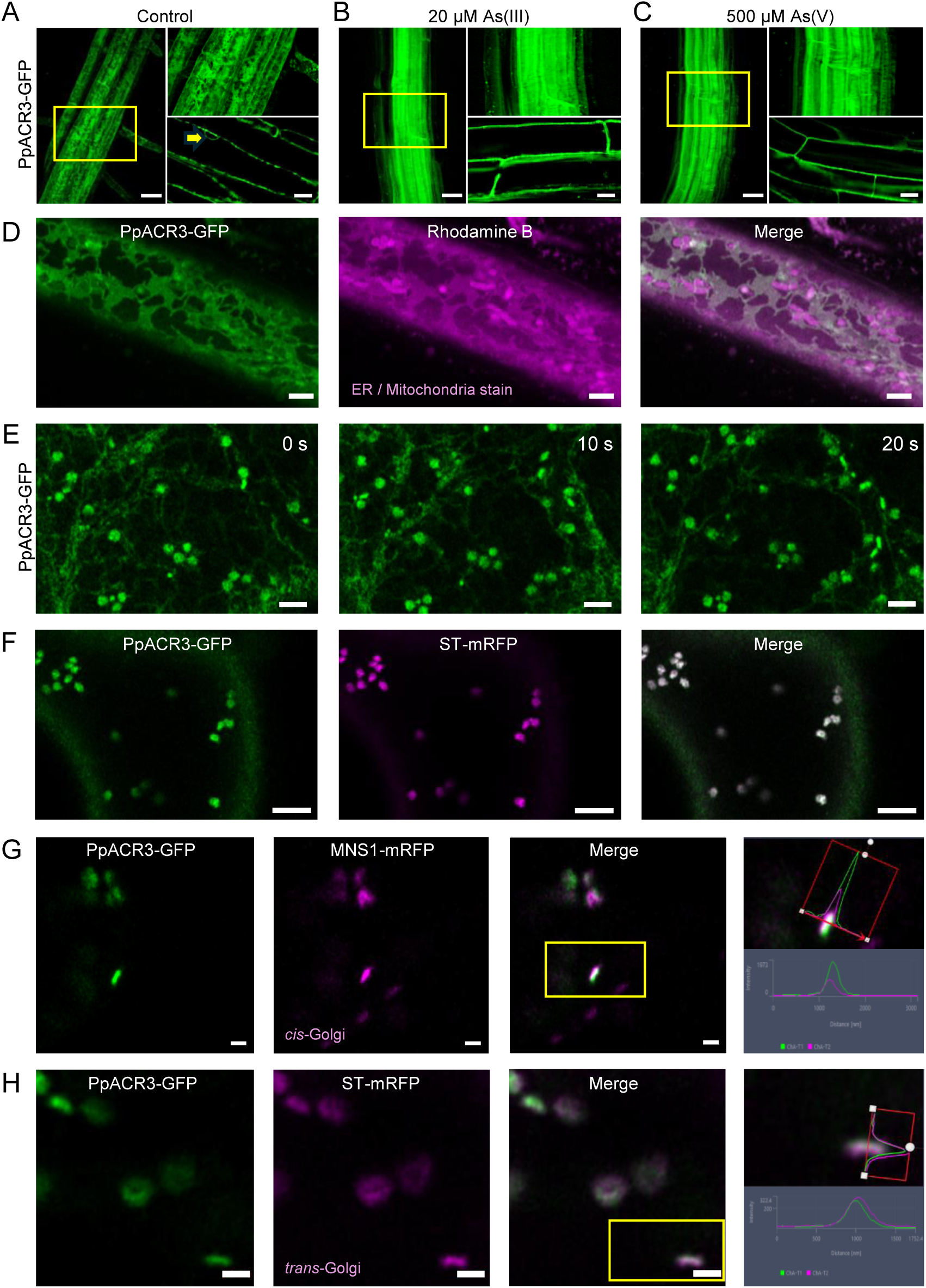
Subcellular localization of PpACR3 in plant cells. (**A**) PpACR3-GFP localized intracellularly in *A. thaliana* root cells (line T1) when grown under basal conditions (MS medium). Yellow arrow indicates the characteristic perinuclear signal. Images for line T10 are shown in Figure S6. (**B, C**) In the presence of As(III) and As(V), PpACR3-GFP localized exclusively to the plasma membrane (PM). PpACR3-GFP expressing seedlings (line T1) were grown for 4 days on MS medium and transferred to MS medium with or without As(III) or As(V) and grown for 24 h before imaging with the use of the ZEISS Lattice Lightsheet 7 microscopy. Roots are displayed either as 3D reconstructions or as 2D views of single planes. Three plants from each treatment group were analyzed, and representative images are shown. Scale bars = 50 µm and 15 µm for 3D and 2 images, respectively. (**D**) Airyscan confocal image of *A. thaliana* root epidermal cells expressing PpACR3-GFP (green) in a stable manner and showing endoplasmic reticulum (ER) localization. Seedlings were stained with rhodamine B hexyl ester (magenta), a lipophilic dye that labels the ER network but also stains mitochondria. Overlay demonstrates substantial co-localization of PpACR3-GFP with rhodamine-labelled ER structures, while discrete punctate magenta signals likely correspond to mitochondria. Scale bars = 2 µm. (**E**) Representative Airyscan confocal time-series images of *A. thaliana* true leaf epidermal cells stably expressing PpACR3-GFP. Images acquired at 0, 10, and 20 s reveal localization of PpACR3-GFP to the ER network and highly mobile punctate structures consistent with Golgi cisternae. Three plants from two independent lines, respectively, were analyzed. Scale bars = 2 µm. (**F-H**) Tobacco leaf epidermal cells transiently co-expressing the indicated proteins with fluorescent tags were visualized by Airyscan high-resolution confocal microscopy 3 days after infiltration. (**F**) Punctate structures of PpACR3-GFP correspond to Golgi bodies labeled with the *trans*-Golgi body marker ST-mRFP (magenta). Scale bars = 5 µm. (**G, H**) Co-localization line profile analysis for PpACR3-GFP with *cis-* and *trans*-Golgi marker constructs. (**G**) Example images and line profile for PpACR3-GFP (green) with the *cis*-Golgi body marker MNS1-mRFP (magenta) and (**H**) with the *trans*-Golgi body marker ST-mRFP (magenta). Line profile applied for analysis and line profile output are shown. Peak distance analysis is shown in Figure S7. Scale bars = 1 µm.

To confirm PpACR3 localization to the ER, *A. thaliana* roots stably expressing PpACR3-GFP were stained with rhodamine B hexyl ester, a lipophilic dye that labels the ER network and mitochondria, and imaged using high-resolution confocal microscopy. Strong co-localization of PpACR3-GFP with rhodamine-labeled ER structures was observed (Figure 5D). Furthermore, PpACR3-GFP also co-localized with the ER network in *A. thaliana* true leaf epidermal cells (Figure 5E).

In addition, PpACR3-GFP signals were detected in highly mobile punctate structures, suggesting localization to Golgi bodies (Figure 5E). To verify this, PpACR3-GFP was transiently co-expressed with Golgi markers in tobacco leaf epidermal cells and analyzed by high-resolution confocal microscopy. PpACR3 localized to punctate structures labeled by the *trans*-Golgi marker ST-mRFP (Figure 5F). Moreover, co-localization line profile analysis showed that PpACR3-GFP did not co-localize with the *cis*-Golgi marker MNS1-mRFP (Figure 5G, Figure S7), but instead co-localized with ST-mRFP, further supporting its localization to *trans*-Golgi cisternae (Figure 5H, Figure S7)

Together, these results indicate that under non-arsenic conditions PpACR3 trafficking to the PM is delayed in plant cells, whereas arsenic exposure promotes redistribution of the transporter from the ER and Golgi to the PM, thereby relieving its intracellular retention.

### 2.7. Conserved N-Terminal Cysteine Residues Control PM Trafficking of Plant ACR3 Transporters

We have previously shown that the ER-to-PM relocalization of MpACR3 in response to arsenicals depends on three arsenic-binding cysteine residues in its arsenic-responsive regulatory domain (Mizio et al. 2025). ACR3 proteins with analogous N-terminal regions (CeACR3, PpACR3, and PsACR3) appear to be regulated similarly (Figure 3) and contain cysteine residues corresponding to those in MpACR3 (Figures 6A, S3). Notably, CeACR3 lacks cysteine residue at the position equivalent to MpACR3 Cys29, whereas Cys68 is not conserved in other ACR3s with elongated N-terminal tails (Figure S3).

**Figure 6.**
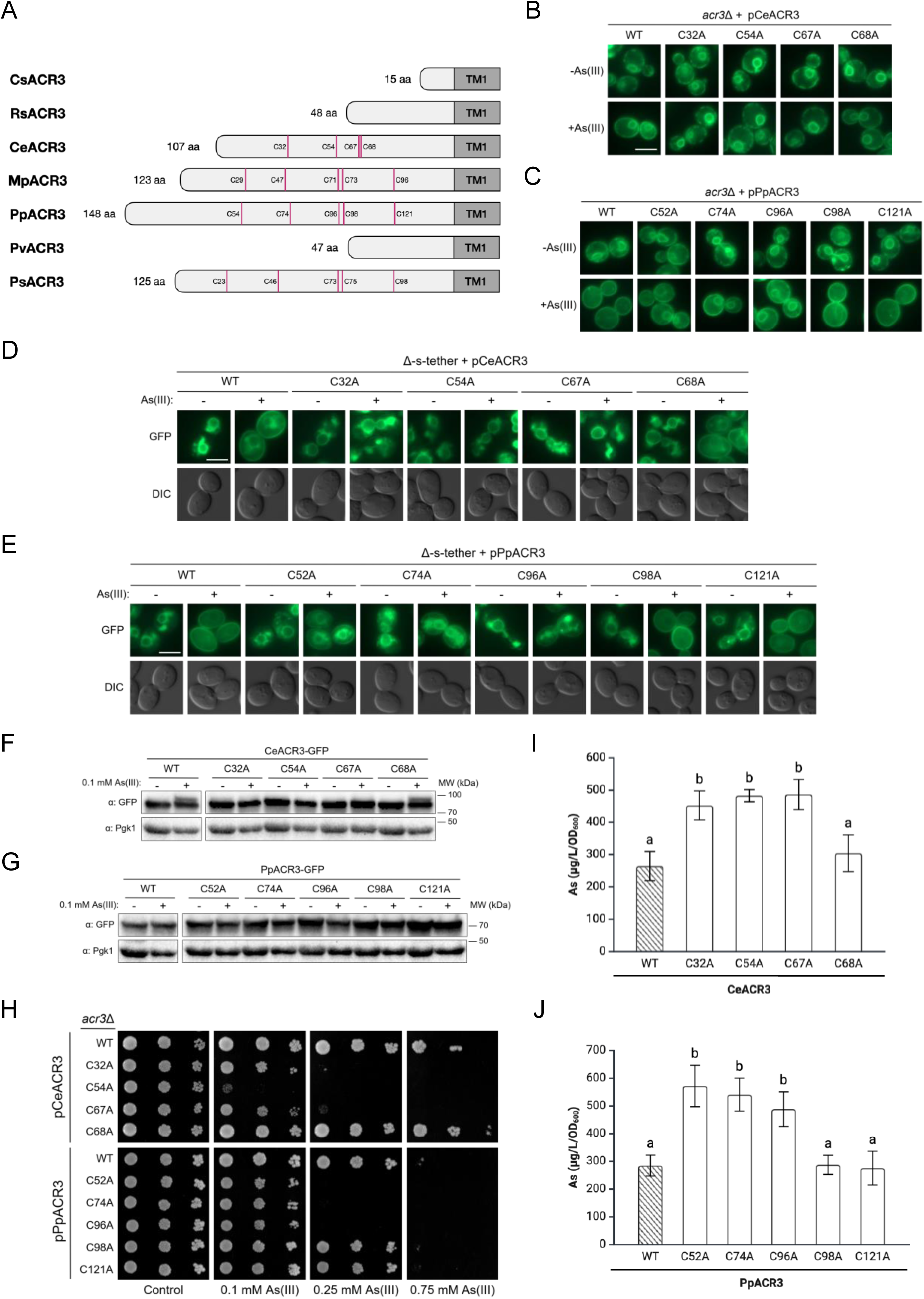
Functional analysis of conserved cysteine residues located in N-terminal tails of CeACR3 and PpACR3. (**A**) Schematic comparison of N-terminal tails of plant ACR3 proteins analyzed in this study. Conserved cysteine residues are highlighted in magenta. TM, transmembrane region. (**B, C**) Subcellular localization of CeACR3, PpACR3, and their mutant variants in *acr3*Δ cells under control conditions or after exposure to 0.1 mM As(III) for 2 h, monitored by fluorescence microscopy. Scale bar = 5 µm. (**D, E**) Localization of the WT and mutated variants of CeACR3 or PpACR3 in Δ-s-tether cells observed under control conditions or following exposure to 0.3 mM As(III) for 2 h by fluorescent microscopy. Scale bar = 5 µm. (**F, G**) Western blot analysis of CeACR3 and PpACR3 variants in *acr3*Δ cells. Yeast cultures were treated with 0.1 mM As(III) or left untreated for 2 h before protein extraction. MW, molecular weight. (**H**) Growth assays of *acr3*Δ cells expressing indicated variants of CeACR3 and PpACR3 with or without As(III). (**I, J**) Arsenic accumulation in *acr3*Δ cells expressing the indicated CeACR3 variants. Yeast cultures were exposed to 1 mM As(III) for 2 h (**I**) or 0.1 mM As(III) for 4 h (**J**) before measurements. Error bars represent mean values ± standard deviation (SD) from three independent experiments (n = 3). Control groups are indicated by diagonal lines. Different letters indicate statistically significant differences determined by one-way ANOVA (p < 0.05).

To test whether these residues have a conserved function and regulate As(III)-dependent trafficking to the PM, we focused on CeACR3 and PpACR3 and replaced individual cysteines with alanine, then assessed localization and function using yeast heterologous system. In CeACR3, substituting any cysteine other than Cys68 caused the fusion proteins to remain in the ER, even in the presence of As(III) (Figure 6B). In PpACR3, alanine substitutions at sites corresponding to Cys29, Cys47, and Cys71 of MpACR3 (i.e., Cys52, Cys74, and Cys96) partially impaired the ER-to-PM relocalization of the fusion proteins after arsenic exposure as the increased PM signal was still observed under these conditions (Figure 6C). In contrast, C98A and C121A variants showed no defect in the ER-to-PM shift (Figure 6C). All above observations were further confirmed in the Δ-s-tether strain (Figure 6D, E).

No overall differences in protein abundance were observed between wild-type and mutant variants; however, mutation of Cys32, Cys54, or Cys67 in CeACR3 abolished the appearance of an additional slowly migrating form upon arsenic exposure (Figure 6F). In contrast, PpACR3 migrated as a single form under all tested conditions (Figure 6G). The identity of the additional CeACR3 form remains to be identified.

Importantly, mutants defective in the arsenic-induced ER-to-PM relocalization also failed to confer wild-type levels of arsenic resistance in *acr3*Δ cells (Figure 6H) and accumulated significantly more arsenic than positive controls (p < 0.05) (Figure 6I, J). In sum, these findings demonstrate that conserved N-terminal cysteine residues are essential for arsenic-responsive trafficking and play a key role in the regulation of plant ACR3 transporters with extended N-termini.

### 2.8. A Conserved Di-Arginine Motif Controls Intracellular Retention of Plant ACR3 Transporters

Our previous research demonstrated that the conserved di-arginine ER-retention motif (L-R-C-R-F) within the arsenic-sensing domain of MpACR3 is essential for trafficking regulation, with the cysteine and the phenylalanine being crucial for the arsenic-induced relocation from the endomembrane system to the PM (Mizio et al. 2025).

To test the functionality of this motif in other plant ACR3 transporters, we generated CeACR3-RR and PpACR3-RR double mutants by substituting the core arginine residues of the motif (Arg31 and Arg33 in CeACR3; Arg73 and Arg75 in PpACR3) with alanine and expressed the resulting variants using the yeast heterologous system (Figure 7A). These variants, together with the analogous MpACR3-RR mutant (Arg46 and Arg48 substituted with alanine), were analyzed for subcellular localization and transport activity. As anticipated, loss of the arginine residues led to strong PM localization of fusion proteins even in the absence of arsenic (Figure 7B), with the result for CeACR3-RR being further confirmed in the Δ-s-tether cells (Figure 7C). Growth assays revealed that disrupting the di-arginine motif significantly impacted arsenic tolerance in *acr3*Δ cells. Specifically, CeACR3-RR expression enhanced yeast growth under arsenic stress, whereas PpACR3-RR and MpACR3-RR reduced arsenic tolerance relative to wild-type proteins (Figure 7D). Consistently, yeast strains expressing mutant ACR3 transporters accumulated decreased (CeACR3-RR) or increased (PpACR3-RR, MpACR3-RR) levels of arsenic compared to controls (p < 0.05) (Figure 7E). These mutations did not affect the overall protein levels, but additional slowly migrating forms of the variant proteins were observed under both control and arsenic stress conditions (Figure 7F).

**Figure 7.**
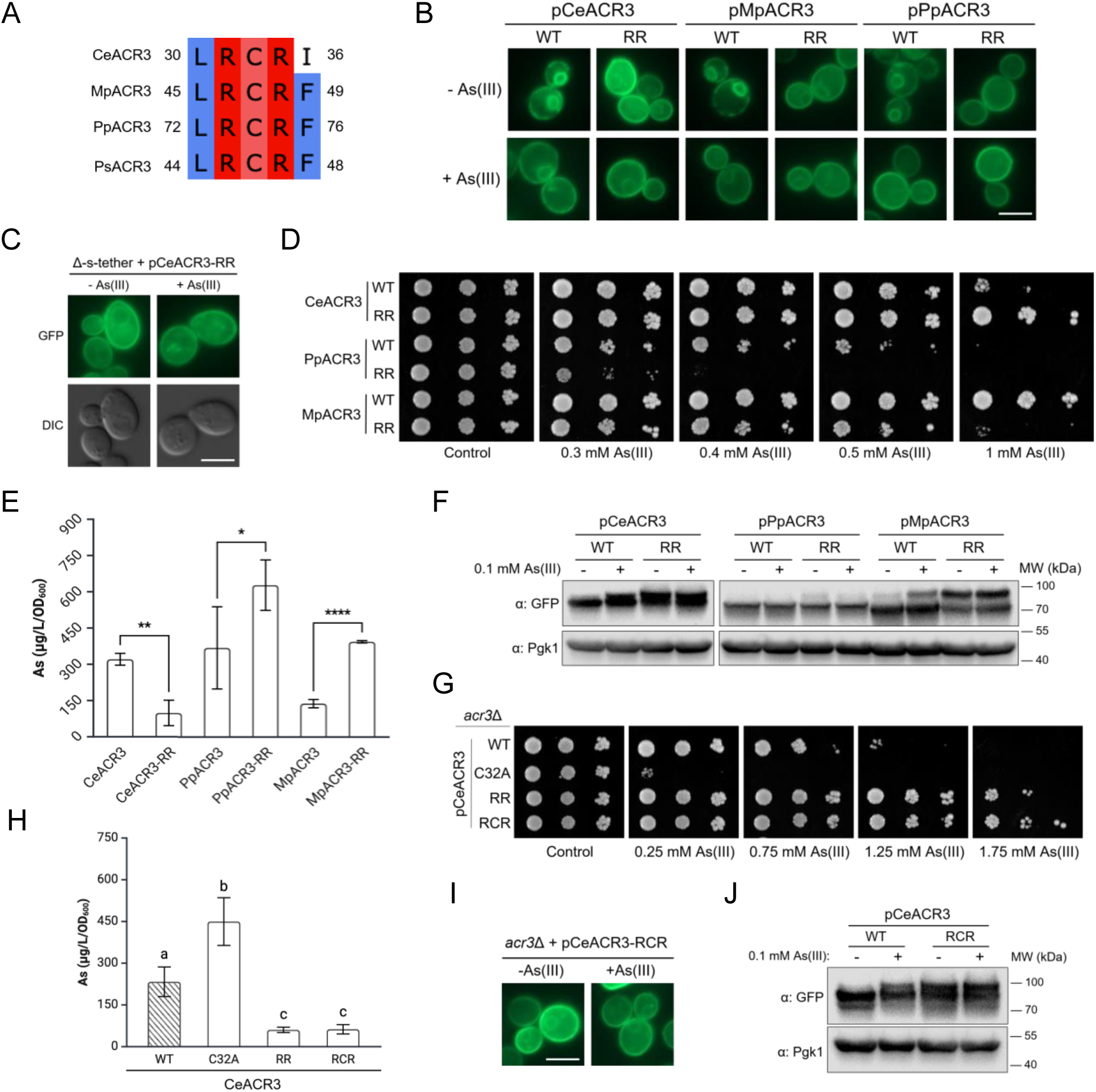
Functional analysis of the di-arginine motif in intracellular retention and function of CeACR3 and PpACR3 in yeast cells. (**A**) Comparison of di-arginine motif sequences in selected ACR3 orthologs based on the multiple sequence alignment presented in Figure S3. (**B**) RR variants of CeACR3, PpACR3, and MpACR3 localize predominantly to the plasma membrane (PM) of *acr3*Δ cells under control conditions and after treatment with 0.1 mM As(III) for 2 h, as determined by fluorescence microscopy. Scale bar = 5 µm. WT, wild-type (**C**) Fluorescence microscopy of Δ-s-tether cells confirms PM localization of CeACR3-RR under the same conditions. Scale bar = 5 µm. (**D**) Growth assays of *acr3*Δ cells expressing WT and RR variants of CeACR3, PpACR3 and MpACR3 under control conditions or in the presence of various concentrations of As(III). (E) Arsenic accumulation in *acr3*Δ cells expressing the indicated ACR3 variants and exposed to 1 mM As(III) for 2 h (CeACR3 and MpACR3) or 0.1 mM As(III) for 4 h (PpACR3). Error bars represent mean values ± standard deviation (SD) from three independent experiments (n = 3). P-values were calculated using a two-tailed *t*-test (*, p < 0.05; **, p < 0.01; ****, p < 0.0001). (F) Western blot analysis of the indicated ACR3 variants in *acr3*Δ cells. Yeast cultures were treated with 0.1 mM As(III) or left untreated for 2 h before protein extraction. MW, molecular weight. (**G**) Growth of *acr3*Δ cells expressing indicated variants of CeACR3 under control conditions or in the presence of various concentrations of As(III). (**H**) Arsenic accumulation in the indicated CeACR3 variants. Error bars represent mean values ± standard deviation (SD) from three independent experiments (n = 3). The control group is indicated by diagonal lines. Different letters indicate statistically significant differences determined by one-way ANOVA (p < 0.05). (**I**) Subcellular localization and (**J**) protein levels of the CeACR3-RCR variant.

Previously, we showed that simultaneous mutation of either arginine residue in the retention motif together with the adjacent Cys47 residue (MpACR3-R46A,C47A and MpACR3-C47A,R48A mutants, respectively) suppresses the arsenic-hypersensitive phenotype and trafficking defect of the MpACR3-C47A single mutant (Mizio et al. 2025). To test whether the arsenic tolerance-enhancing RR mutation of CeACR3 would also rescue the CeACR3-C32A phenotype, we constructed a triple variant with Arg31, Cys32 and Arg33 residues of CeACR3 substituted for alanine (CeACR3-RCR) and analyzed it with the wild-type, C32A and RR variants of CeACR3. Indeed, the RR mutation remained dominant over the C32A mutation, as evidenced by comparable growth on solid media (Figure 7G), arsenic accumulation (Figure 7H), and subcellular localization (Figures 6B, 7B, I) observed in the *acr3*Δ cells expressing CeACR3-RR and CeACR3-RCR variants. Importantly, the protein level of the CeACR3-RCR was not affected compared to WT or other variants (CeACR3-C32A and CeACR3-RR) (Figures 6F, G, 7F, J). Taken together, these results confirm the role of the conserved di-arginine motif as a pivotal regulator of plant ACR3 trafficking.

## 3. Discussion

Arsenic stress has acted as a strong evolutionary pressure, driving the emergence of specialized detoxification mechanisms as early as in anoxic organisms predating the Great Oxidation Event (Chen et al. 2020). Across taxa, from prokaryotes to fungi, extrusion of As(III) mediated by ACR3 transporter orthologs represents a major detoxification strategy, conferring tolerance to millimolar concentrations of this metalloid (Maciaszczyk-Dziubinska et al. 2012; Garbinski et al. 2019). Although genes encoding ACR3 orthologs are also widespread in plants, their role in metalloid detoxification remains poorly understood, despite the ongoing global threat of arsenic contamination.

To date, research on plant ACR3 transporters has been largely focused on the arsenic-hyperaccumulating fern *P. vittata* and its close relatives. The presence of multiple ACR3 paralogs, exhibiting diverse subcellular localization patterns, including the PM and VM, makes these species valuable models for phytoremediation studies (Danh et al. 2014). Moreover, it has been demonstrated in *P. vittata* that As(V) is metabolized to 1-arseno-3-phosphoglycerate, which is subsequently accumulated in intracellular vesicles, followed by the release of As(V) and its reduction to As(III) (Cai et al. 2019). However, these systems appear to be restricted to this genus and may not be representative of detoxification mechanisms in other ACR3-containing plants.

In this study, we used the well-established eukaryotic model *S. cerevisiae* lacking the endogenous *ACR3* gene to functionally characterize ACR3 transporters from plant species representing evolutionarily distant phylogenetic lineages, including green algae (CsACR3 from *C. subellipsoidea*, RsACR3 from *R. subcapitata*, and CeACR3 from *C. eustigma*), bryophytes (PpACR3 from *P. patens*), and gymnosperms (PsACR3 from *P. sitchensis*) (Figure 1). The ecological diversity of the selected organisms (Table S2), together with the inclusion of previously characterized MpACR3 from the liverwort *M. polymorpha* (Li et al. 2024; Dutta et al. 2024; Mizio et al. 2025), and PvACR3 from the fern *P. vittata* (Chen et al. 2017; Chen et al. 2021) as reference transporters, provided a robust framework for comparative functional analysis of ACR3 proteins.

By combining functional assays with live-cell imaging in *S. cerevisiae*, we demonstrated that all tested ACR3 proteins are functional PM transporters mediating As(III) efflux (Figure 3). These results are consistent with our predicted topology of plant ACR3 proteins, featuring 10 transmembrane regions with both the N- and C-termini oriented toward the cytoplasm (Figure S2). Such orientation is critical for As(III) extrusion, as transport is driven by the proton motive force across the PM (Maciaszczyk-Dziubińska et al. 2011; Villadangos et al. 2012). In addition, the C-terminal GFP fusion yielded a strong fluorescence signal in the PM, which would be substantially reduced under the acidic conditions used in this study (pH < 6) if localized extracellularly (Materials and Methods; Figure 3) (Alkaabi et al. 2005).

The tested plant ACR3 orthologs conferred differential arsenic tolerance in the *acr3*Δ yeast mutant, with some proteins enhancing resistance to both As(III) and As(V) (CsACR3, CeACR3, MpACR3, and PvACR3), whereas others substantially increased tolerance to As(V) but conferred only moderate (PpACR3) or weak (RsACR3 and PsACR3) resistance to As(III) (Figure 3). These differences are unlikely to reflect intrinsic substrate specificity, as As(V) is rapidly reduced to As(III) by the arsenate reductase Acr2 in yeast, ultimately generating a common intracellular As(III) pool that is subsequently extruded by Acr3 (Mukhopadhyay and Rosen 1998). Instead, the observed variation likely reflects the heterologous context of *S. cerevisiae*, where folding and trafficking limitations may contribute to the differential tolerance phenotypes, as ACR3 activity depends on its efficient accumulation at the PM. Even subtle differences in targeting efficiency to the PM could substantially influence transport capacity in yeast cells (Figure 3). This is particularly evident for PsACR3, which exhibited pronounced intracellular retention and the lowest level of As(III) tolerance (Figures 2, 3). Furthermore, differences in membrane lipid composition and sterol content between yeast and plant cells may influence ACR3 activity, as membrane physical properties directly affect transporter folding, conformational dynamics, and functional regulation (Ernst and Robertson 2021; Stieger et al. 2021). Consistent with this interpretation, PpACR3 confers comparable tolerance to both As(III) and As(V) when expressed in *A. thaliana* (Figure 4), in contrast to its more selective effect in yeast (Figure 3). Moreover, apparent folding defects leading to vacuolar degradation precluded reliable functional characterization of ACR3 from the filamentous alga *Klebsormidium nitens* (KnACR3) and the lycophyte *S. moellendorffii* (SmACR3) (data not shown).

Together, these findings suggest that the differential tolerance observed in yeast primarily reflects context-dependent regulation, trafficking, and membrane environment rather than inherent functional divergence among plant ACR3 proteins. Consequently, accurate assessment of ACR3 transport properties will require analysis in native systems or plant-based models.

ACR3 proteins are generally thought to be regulated at the transcriptional level (Maciaszczyk-Dziubinska et al. 2012; Garbinski et al. 2019), with limited information available on post-translational mechanisms controlling their activity. We have recently shown that MpACR3 from the liverwort *M. polymorpha* displays dual localization to the PM and endomembrane compartments, suggesting a more complex regulatory behavior (Mizio et al. 2025). In both *M. polymorpha* and *A. thaliana,* MpACR3 predominantly associates with the Golgi network, whereas in yeast it is retained in the ER. Notably, its redistribution from the Golgi/ER to the PM in response to arsenic points to a dynamic, stimulus-dependent trafficking mechanism (Mizio et al. 2025). This regulation appears to be mediated by an extended N-terminal cytosolic metalloid-sensing domain that coordinates arsenic via three conserved cysteine residues, including one embedded within a di-arginine motif implicated in intracellular retention (Michelsen et al. 2005; Banfield 2011; Lujan and Campelo 2021; Mizio et al. 2025).

Here, we show that arsenic-driven sorting to the PM is a conserved feature of plant ACR3 proteins possessing an extended cysteine-rich N-terminus (CeACR3, MpACR3, PpACR3, and PsACR3) (Figure 3; Figures S1-S3). In the absence of arsenic, these proteins localized predominantly to the endomembrane system (Figures 3, 5). Among these, cortical and perinuclear ER were identified in yeast cells. Strikingly, arsenic exposure promoted relocalization of tested ACR3-GFP proteins to the PM, with residual (MpACR3 and PpACR3), moderate (CeACR3), or substantial (PsACR3) signal remaining in the endomembrane system (Figure 3B, D). Importantly, when expressed in *A. thaliana*, PpACR3 localized to both the ER/Golgi and PM under control conditions (Figure 5), whereas arsenic treatment resulted in its accumulation predominantly at the PM (Figure 5B, C). These observations support the notion that the behavior of plant ACR3 proteins with extended cysteine-rich N-termini in yeast reflects their subcellular localization patterns and arsenic-dependent regulation in their native systems. However, the subcellular localization patterns of PpACR3 in the absence and presence of arsenic need to be verified in its native organism.

Next, we sought to identify N-terminal cysteine residues involved in arsenic-induced relocalization from the ER/Golgi to the PM, as previously demonstrated for MpACR3 (Mizio et al. 2025). For mutational analysis, we selected PpACR3, which contains five conserved cysteine residues in its N-terminus and (Figures 6, S1, S3) and exhibits arsenic-driven PM trafficking in both yeast and plants (Figures 3, 5). In addition, we generated Cys-to-Ala variants of CeACR3, which contains only four cysteine residues and lacks the first N-terminal cysteine residue located upstream of the L-R-C-R-F motif and shared among plant ACR3 proteins, including PpACR3 (Figures S1 and S3). Importantly, this residue was previously shown to participate in arsenic sensing and to promote MpACR3 accumulation at the PM (Mizio et al. 2025).

For each protein, we identified three cysteine residues required for efficient accumulation at the PM and, consequently, for conferring arsenic tolerance (Figure 6). This finding is consistent with the proposed role of the extended N-terminus as an arsenic-sensing module that coordinates As(III) via three cysteine residues (Mizio et al. 2025). The observation that CeACR3 utilizes a distinct set of N-terminal cysteine residues to mediate arsenic-driven relocalization from the endomembrane system to the PM suggests that, despite the low sequence identity of cysteine-rich N-terminal domains among plant ACR3 proteins (Figure S3), these domains may share a conserved mechanism of arsenic sensing and regulated PM trafficking.

We previously proposed that As(III) binding to a predicted N-terminal β-barrel–like domain of MpACR3 induces a conformational change that promotes release from ER or Golgi retention, likely through modulation of an N-terminal di-arginine motif that is essential for intracellular localization in the absence of arsenic (Mizio et al. 2025). Based on sequence alignments and alanine-scanning mutagenesis, we defined a consensus motif, Φ₁-R-C-R-Φ₂ (where Φ denotes a bulky hydrophobic residue), that regulates intracellular retention and arsenic-driven relocalization to the PM. Furthermore, we found that the cysteine and Φ_2_ residues within this motif are required for arsenic-induced trafficking, whereas the Φ_1_ and two arginine residues contribute primarily to intracellular retention (Mizio et al. 2025).

Our structural predictions indicate that, despite low sequence similarity, the elongated N-termini of all tested ACR3 proteins adopt similar β-barrel-like folds (Figure S8) containing a highly conserved L-R-C-R-F/I motif (Figures S1, S3). Furthermore, substitution of conserved arginine residues within this motif in CeACR3 and PpACR3 resulted in exclusive PM localization, supporting their role in retaining ACR3 in intracellular compartments in the absence of arsenic (Figure 7). Interestingly, both MpACR3-RR and PpACR3-RR variants lacking di-arginine motif exhibited reduced functionality despite their predominant localization at the PM, whereas expression of the CeACR3-RR variant conferred increased tolerance to As(III) (Figure 7). Thus, the conserved RR residues primarily mediate intracellular retention, but their divergent functional effects suggest that RR motif also influences the conformation and activity of the N-terminal regulatory domain. This motif could therefore be viewed as part of an arsenic-responsive module coupling retention, conformational control, and transporter function. Predicted structural models of the N-terminal domains of MpACR3-RR and PpACR3-RR suggest a more relaxed architecture lacking defined α-helical elements (Figures S9, S10). In contrast, substitution of arginine residues in CeACR3 resulted in a highly disordered N-terminal structure that may not interfere with the PM domain of CeACR3 (Figure S11). This is consistent with the increased tolerance to arsenic conferred by the expression of CeACR3-RR variant (Figure 7). Whether these findings reflect a broader role of N-terminal conformational dynamics in regulating ACR3 activity at the PM remains to be determined and will require direct measurements of transport activity in the corresponding plant ACR3 variants.

In addition to arsenic-driven trafficking to the PM, increased accumulation of MpACR3 in both *M. polymorpha* and *A. thaliana* (Mizio et al. 2025) and PpACR3-GFP in *A. thaliana* (Figure 4A) was observed upon arsenic treatment. In the case of MpACR3, this likely results from transcriptional upregulation and increased MpACR3 mRNA accumulation (Mizio et al. 2025). In contrast, PpACR3 transcript levels were not elevated following arsenic treatment in its native organism (Figure S4), suggesting that enhanced protein stability and/or reduced degradation may also contribute to the regulation of plant ACR3 proteins. These observations warrant further investigation to elucidate the molecular mechanisms underlying transcriptional and post-translational regulation of plant ACR3 proteins.

Understanding the evolutionary diversity and regulation of ACR3 arsenite transporters provides critical insight into natural mechanisms of arsenic detoxification operating across diverse plant lineages. By identifying both constitutive and metalloid-responsive arsenic efflux systems, this work highlights molecular features that could be leveraged to improve phytoremediation strategies in arsenic-contaminated environments. Moreover, the discovery of metalloid-sensitive regulatory domains may provide a promising foundation for the development of highly sensitive biological tools for arsenic detection, although further engineering and validation will be required before their potential application in environmental monitoring and hazard assessment.

## 4. Materials and Methods

### 4.1. Yeast Strains, Growth Conditions, and Transformation

The *S. cerevisiae* strains used in this study included RW104 (*acr3*Δ*::kanMX6*) (Wysocki et al. 2001) and RW104 expressing *TRP1*::*proTPI1-SS-mCherry-HDEL* in the W303-1A background (*MAT*a *leu2-3,112 trp1-1 can1-100 ura3-1 ade2-1 his3-11,15*) (Mizio et al. 2025), as well as Δ-s-tether (*ist2*Δ*::hisMX6 scs2*Δ*::TRP1 scs22*Δ*::hisMX6 tcb1*Δ*::kanMX6 tcb2*Δ*::kanMX6 tcb3*Δ*::hisMX6 ice2*Δ*::natMX4*) in the SEY6210 background (*MAT*α *leu2-3,112 ura3-52 his3-*Δ*200 trp1-*Δ*901 suc2-*Δ*9 lys2-801; GAL*) (Quon et al. 2018). The Δ-s-tether strain was kindly provided by Christopher T. Beh (Simon Fraser University, British Columbia, Canada). Yeast cells were grown at 30°C in standard synthetic dextrose (SD) medium (pH 5.4) supplemented as required.

To assess sensitivity to arsenic compounds, mid-log phase cultures were subjected to 10-fold serial dilutions and spotted onto solid SD medium containing the indicated concentrations of sodium arsenite [As(III)] or sodium arsenate [As(V)] (Sigma-Aldrich). Plates were imaged after 3 days of incubation at 30°C. For subcellular localization analyzes, mid-log phase cultures expressing the indicated fusion proteins were incubated with the specified concentrations of metalloids for 2 h prior to microscopy. Yeast transformation was performed using the lithium acetate/single-stranded carrier DNA/polyethylene glycol method (Gietz and Woods 2002).

### 4.2. Plant Material and Growth Conditions

Plants of *A. thaliana* ecotype Col-0 and transgenic lines expressing *pro35S:PpACR3-GFP* were grown under a 16 h light/8 h dark photoperiod at 22°C. Cultivation was performed either in soil or in vitro on half-strength Murashige and Skoog medium (MS; Sigma-Aldrich) supplemented with 1% (w/v) sucrose. For phenotypic comparisons, wild-type and *pro35S:PpACR3-GFP* transgenic lines (T1 and T10) were cultivated in soil. At the end of the growth cycle, the following parameters were recorded: primary inflorescence height, number of flowers on the main stem, total branch number, and total flower number.

For germination assays, surface-sterilized seeds from wild-type and transgenic lines were sown on MS medium supplemented with either 50 µM As(III) or 300 µM As(V), or on control medium lacking arsenic, and grown for 14 days.

For seedling growth assays, seeds were first germinated on arsenic-free MS medium. After 4 days, seedlings were transferred to MS medium containing 20 µM As(III), 500 µM As(V), or no arsenic (control), and cultivated for an additional 7 days. For microscopy analyses, seedlings were initially grown on MS medium without arsenic. After 4 days, they were transferred to MS medium supplemented with 25 µM As(III), 1000 µM As(V), or no arsenic, and incubated for a further 24 h prior to imaging.

The wild-type *P. patens* ecotype Gransden 2004 (kindly provided by Piotr Wasko, University of Maria Curie-Sklodowska) was grown under long day (LD,16 h light/8 h dark) photoperiod and 22°C on solid BCD medium (1 mM CaCl_2_, 45 µM FeSO_4_•7H_2_O, 1 mM MgSO_4_, 1.84 mM KH_2_PO_4_, 10 mM KNO_3_, Hoagland’s A-Z trace element solution) and 0.8% (w/v) agar (A4800, Sigma-Aldrich). For *ACR3* expression assays, plants were cultured on BCD medium supplemented with As(III) or As(V) at indicated concentrations.

### 4.3. Plasmids and Gene Cloning

Oligonucleotides used for ACR3 cloning are listed in Table S5, and plasmids are summarized in Table S6. Coding sequences of *CsACR3, RsACR3*, *CeACR3*, *PpACR3*, *PvACR3*, and *PsACR3* were synthesized (Thermo Fisher Scientific) based on sequences retrieved from the NCBI database. The genes were cloned into the pUG35 vector as C-terminal GFP fusions under the control of the constitutive *MET17* promoter using the In-Fusion HD Cloning Kit (Takara). All constructs (pCsACR3, pRsACR3, pCeACR3, pPpACR3, pPvACR3, and pPsACR3) were verified by DNA sequencing. The pMpACR3 construct was described previously (Mizio et al. 2025).

For expression in plants, the synthetic *PpACR3* gene was amplified by PCR (PrimeSTAR® Max DNA Polymerase, Takara Bio) and cloned (In-Fusion® HD Cloning Kit, Takara Bio) into the pGFPGUSPlus plasmid (Vickers et al. 2007) (a gift from Claudia Vickers; Addgene plasmid #64401; http://n2t.net/addgene:64401) under the control of the constitutive *35S* promoter in fusion with the C-terminal GFP tag-coding sequence. The obtained p35S::PpACR3-GFP plasmid was confirmed by DNA sequencing. The oligonucleotides used for cloning and sequencing are listed in Table S5.

### 4.4. Mutagenesis

Plasmids containing wild-type *ACR3-GFP* fusion genes were used as a template for site-directed mutagenesis employing QuikChange Lightning kit (Agilent Technologies). Mutations were confirmed by DNA sequencing. Oligonucleotides used for mutagenesis and sequencing are listed in Table S5.

### 4.5. Plant Transformation

The *pro35S:PpACR3-GFP* construct was introduced into *Agrobacterium tumefaciens* strain GV3101 (pMP90) (Koncz and Schell 1986) by electroporation and subsequently used to transform *A. thaliana* (Col-0) plants via the floral dip method (Clough and Bent 1998). Seeds from transformed plants were surface-sterilized and sown on MS medium supplemented with 25 μg mL⁻¹ hygromycin B (Sigma-Aldrich). Following 6 h of light exposure, seeds were incubated in darkness for 4 days and then transferred to a 16 h light/8 h dark photoperiod for further growth. Transgenic seedlings were selected based on elongated hypocotyls and the presence of green true leaves and subsequently transferred to soil. T1 and T10 primary transformant lines, exhibiting the strongest pro35S:PpACR3-GFP fluorescence, were selected for further analysis.

### 4.6. Quantitative Real-Time PCR (qRT-PCR)

Total RNA was isolated from 50 mg of plant tissue using the Beadbeat Total RNA Mini Kit (A&A Biotechnology, Gdansk, Poland). First-strand cDNA was synthesized from RNA samples using the High-Capacity cDNA Reverse Transcription Kit (Applied Biosystems). qRT-PCR was performed with cDNA as template using the 2× PCR Master Mix SYBR kit (A&A Biotechnology) on a CFX Connect Real-Time PCR Detection System (Bio-Rad, Hercules, CA, USA) in a final reaction volume of 15 μL. Primer sequences are listed in Table S5. The qPCR cycling conditions were as follows: 1 min at 95°C, followed by 40 cycles of 10 s at 95°C, 15 s at 60°C, and 20 s at 72°C. A melting curve analysis was performed to confirm amplification specificity. Relative gene expression levels were calculated using the comparative Ct (ΔΔCt) method.

### 4.7. Microscopy

Subcellular localization of ACR3–GFP fusion proteins in live *S. cerevisiae* cells was examined using an upright wide-field epifluorescence microscope (Axio Imager M2; Carl Zeiss, Germany) equipped with an HBO 100 illuminator, a 100× oil immersion objective (Plan-Neofluar 100×/1.30), and filter sets 38 HE (eGFP) and 43 HE (mCherry). Images were acquired with a Zeiss AxioCam MRc digital camera and processed using ZEN 3.7 software (Zeiss).

For imaging in *A. thaliana* root cells, a Lattice Light Sheet 7 (LLS7; Carl Zeiss) microscope was used to visualize PpACR3-GFP (488 nm excitation). Imaging parameters were as follows: Sinc3 30 × 1000, z-step of 0.3 µm, 488 nm laser power at 6%, and exposure time of 50 ms. Image processing was performed using ZEN 3.10 (Carl Zeiss Microscopy GmbH), including medium deconvolution, linear interpolation, deskewing, cover glass transformation, and subset selection. Final renderings were generated as 3D projections.

Rhodamine B hexyl ester was used as a lipophilic dye to label the ER in *A. thaliana* roots expressing PpACR3-GFP. Ten-day-old seedlings were incubated in a 1 µM working solution of rhodamine B hexyl ester for 15 min, followed by washing in water to remove excess dye. Imaging was performed using a Zeiss LSM880 confocal microscope with an AiryScan detector and 63×/1.4 NA oil objective. For co-localization analysis with PpACR3-GFP, dual-channel imaging was employed. GFP was excited at 488 nm, and emission was collected between 495 and 550 nm. Rhodamine B hexyl ester was excited at 561 nm, with emission detected between 570 and 615 nm. Images were acquired in line-switching mode with 4× line averaging.

Co-localization of MpACR3-GFP with the marker proteins of Golgi compartments in tobacco leaf epidermal cells was visualized using Airyscan high-resolution confocal microscopy 3 d post-infiltration (Figure 5). The analysis methodology was based on (McGinness et al. 2022; McGinness et al. 2025).

### 4.8. Metalloid Accumulation Measurements

Exponentially growing *S*. *cerevisiae* cultures were treated with the indicated concentrations of As(III) for the specified time, after which cell density was determined by measuring OD_600_. Cells were then harvested by centrifugation, washed three times with ice-cold deionized water, and lysed in 2 mL of 0.5% nitric acid (EMPLURA, Merck) by boiling for 10 min. Following centrifugation (10 min, 4°C), arsenic concentrations in the supernatants were quantified using atomic absorption spectrometry with a graphite furnace (contrAA 800 G, Analytik Jena).

For plant material, a comparable protocol was applied. *A. thaliana* plants were cultivated on solid Murashige and Skoog medium (MS) or 0.5× Gamborg B5 medium supplemented with the indicated concentrations of As(III) or As(V) for 14 or 21 days. Tissue samples were weighed, snap-frozen in liquid nitrogen, and homogenized in 1.5 mL of 0.5% nitric acid using glass beads and a Precellys 24 homogenizer (Bertin Technologies). Homogenates were boiled and centrifuged, and metalloid concentrations in the resulting supernatants were determined as described above.

### 4.9. Protein Extraction and Immunoblotting

Total protein extracts from yeast cells were obtained using the trichloroacetic acid (TCA) method (Wright et al. 1989). For *A. thaliana*, total protein extracts were obtained following the protocol of Martínez-García et al. (1999). Protein extracts were separated by 10% SDS-PAGE, blotted onto nitrocellulose membranes (Bio-Rad), and incubated with anti-GFP antibody (Roche, 11814460001; 1:3000 dilution) for detection of the GFP-tagged ACR3 proteins. Staining nitrocellulose membrane with Ponceau S solution (Sigma-Aldrich) was used for total protein normalization. An additional loading control was performed by the detection of 3-phosphoglycerate (Pgk1) using anti-Pgk1 antibody (Abcam, 22C5D8; 1:5000 dilution).

### 4.10. Phylogenetic Tree, Sequence Alignments and Topology Prediction

The ACR3 protein sequences were downloaded from the following databases: www.ncbi.nlm.nih.gov, www.gigadb.org, www.fernbase.org, www.hornworts.uzh.ch, datadryad.org, phytozome-next.jgi.doe.gov, phycocosm.jgi.doe.gov. The sequence details are provided in Table S1. To construct the phylogenetic tree of plant ACR3 proteins, sequences were aligned using ClustalW 2.1 implemented in Geneious Prime (version 2019.1.1; Biomatters Ltd.) with default settings. Phylogenetic relationships were inferred using the maximum likelihood method implemented on the IQ-TREE web server (Trifinopoulos et al. 2016). Branch support values were estimated using 1000 bootstrap replicates. The resulting phylogenetic tree was visualized using Interactive Tree Of Life (iTOL v3) (Letunic and Bork 2019).

Multiple sequence alignment of plant ACR3 orthologs experimentally analyzed in this study was performed using the Clustal Omega algorithm implemented in Unipro UGENE software (Okonechnikov et al. 2012) with default settings and edited using JalView Version 2 (Waterhouse et al. 2009). Pairwise sequence identity and similarity were calculated using SIAS (http://imed.med.ucm.es/Tools/sias.html) with default parameters. Multiple sequence alignments of short and extended N-terminal ACR3 regions were generated using the Clustal Omega server (https://www.ebi.ac.uk/jdispatcher/msa/clustalo) with default settings and manually adjusted. Membrane topology was predicted using the CCTOP server (https://cctop.ttk.hu) and visualized with Protter 1.0 (https://wlab.ethz.ch/protter).

### 4.11. Protein Structure Prediction

Protein structures were predicted using AlphaFold Server 3 (https://alphafoldserver.com) (Abramson et al. 2024). For each protein sequence, a set of 20 predefined random seeds was used to generate multiple structural models, allowing assessment of prediction variability and consistency. Predicted models were ranked according to the predicted Template Modeling (pTM) score, and only models with pTM ≥ 0.7 were retained for further analysis. When multiple models for a given protein met this threshold, the model with the highest predicted Local Distance Difference Test (pLDDT) score, calculated as the average per-residue confidence score among the top-ranked pTM models, was selected as the most reliable structure. Models of RR variants were generated using the seed numbers corresponding to the highest-scoring wild-type models. Scores and seed numbers for all analyzed models are provided in Table S7. Structures were visualized and analyzed using UCSF ChimeraX 1.9 (Meng et al. 2023).

## Acknowledgements

This work was financially supported by the National Science Centre, Poland (grant no. 2019/35/B/NZ3/00379) awarded to R.W. The authors are grateful to Christopher T. Beh (Simon Fraser University) for generously providing the Δ-s-tether strain and Piotr Wasko (University of Maria Curie-Sklodowska) for the wild-type *P. patens* ssp. *patens* (Gransden 2004 ecotype). LL7 studies were performed at the Laboratory of Lattice Lightsheet Microscopy, Faculty of Biological Sciences, University of Wroclaw.

## Data Availability Statement

Structure prediction data have been deposited in the ModelArchive (https://www.modelarchive.org) database with the accession codes: CeACR3 (ma-71r2v), MpACR3 (ma-yoefs), PpACR3 (ma-33rgr), PsACR3 (ma-0aa6h), CeACR3-RR (ma-rjyal), MpACR3-RR (ma-ggpxu), and PpACR3-RR (ma-d8fyt).

## Supporting Information

**Figure S1.** Multiple sequence alignment of plant ACR3 proteins characterized in this study.

**Figure S2.** Plant ACR3 orthologs share a 10-transmembrane-region topology with both termini oriented toward the cytoplasm.

**Figure S3.** Multiple sequence alignment of N-terminal regions of plant ACR3 proteins analyzed *in silico*.

**Figure S4.** Arsenic exposure does not affect *PpACR3* expression.

**Figure S5.** No phenotypic differences were observed between wild-type (WT, Col-0) *Arabidopsis thaliana* plants and pro35S:PpACR3-GFP transgenic lines (T1 and T10) under control conditions.

**Figure S6.** Subcellular localization of PpACR3 in plant cells.

**Figure S7.** Co-localization line profile analysis for PpACR3-GFP with *cis*- and *trans*-Golgi marker constructs.

**Figure S8.** Comparison of predicted N-terminal tail structures of selected plant ACR3 orthologs.

**Figure S9.** Comparison of predicted N-terminal domain structures of the WT and RR variants of MpACR3.

**Figure S10.** Comparison of predicted N-terminal domain structures of the WT and RR variants of PpACR3.

**Figure S11.** Comparison of predicted N-terminal domain structures of the WT and RR variants of CeACR3.

**Table S1.** Plant ACR3 proteins analyzed *in silico* in this study.

**Table S2.** Characteristics of the plant species selected for this study.

**Table S3.** Pairwise sequence identity and similarity among plant ACR3 orthologs.

**Table S4.** Predicted topology of plant ACR3 transporters.

**Table S5.** Oligonucleotides used in this work.

**Table S6.** Plasmids used in this work.

**Table S7.** Model scores and seed numbers for the N-terminal domains of the indicated ACR3 proteins.

