## Supplementary Materials for "Arsenic-Driven Trafficking of ACR3 Transporters Is Conserved across Plant Lineages"

### Supplementary Figures S1-S11

### Supplementary Tables S1-S7

### Supplementary references

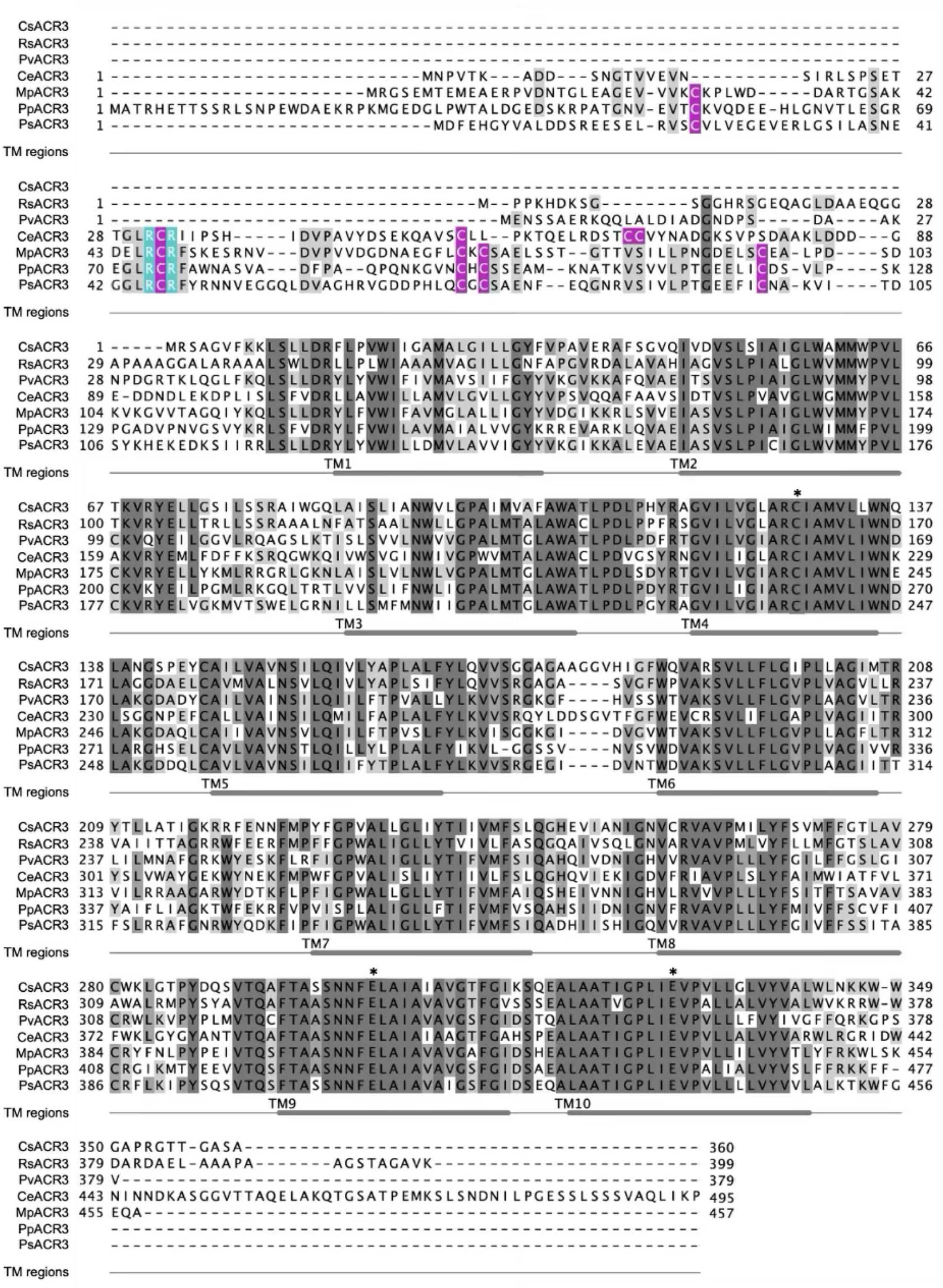

**Figure S1. Multiple sequence alignment of plant ACR3 proteins characterized in this study**. At least 30% sequence identity in a column is highlighted. N-terminal cysteine residues are indicated in magenta. The conserved di-arginine motif is highlighted in cyan. Conserved residues potentially involved in metalloid transport are marked with asterisk. Transmembrane regions (TM) were predicted using CCTOP server (https://cctop.ttk.hu) and represented by grey bars based on representative MpACR3 topology. The alignment was performed using the Clustal Omega algorithm in the Unipro UGENE software (Okonechnikov et al., 2012) and edited using the JalView Version 2 software (Waterhouse et al., 2009). Sequence details are provided in Table S1.

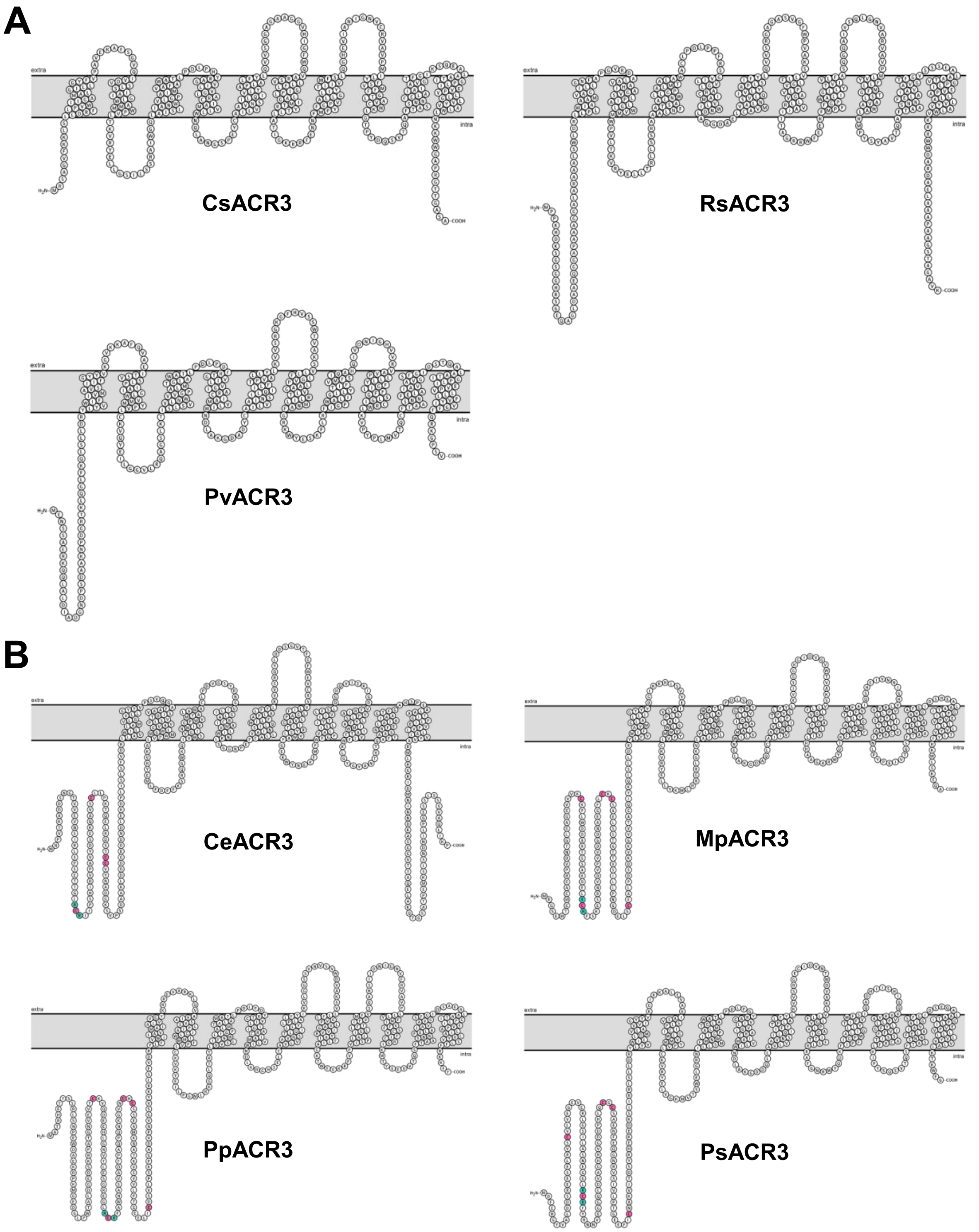

**Figure S2. Plant ACR3 orthologs share a 10-transmembrane-region topology with both termini oriented toward the cytoplasm.** (A, B) Schematic representation of the predicted topology of short-tailed (A) and long-tailed (B) plant ACR3 proteins. N-terminal cysteine residues are highlighted in magenta, and the conserved di-arginine motif is indicated in cyan. The schemes were generated using Protter 1.0 (https://wlab.ethz.ch/protter) and manually adjusted.

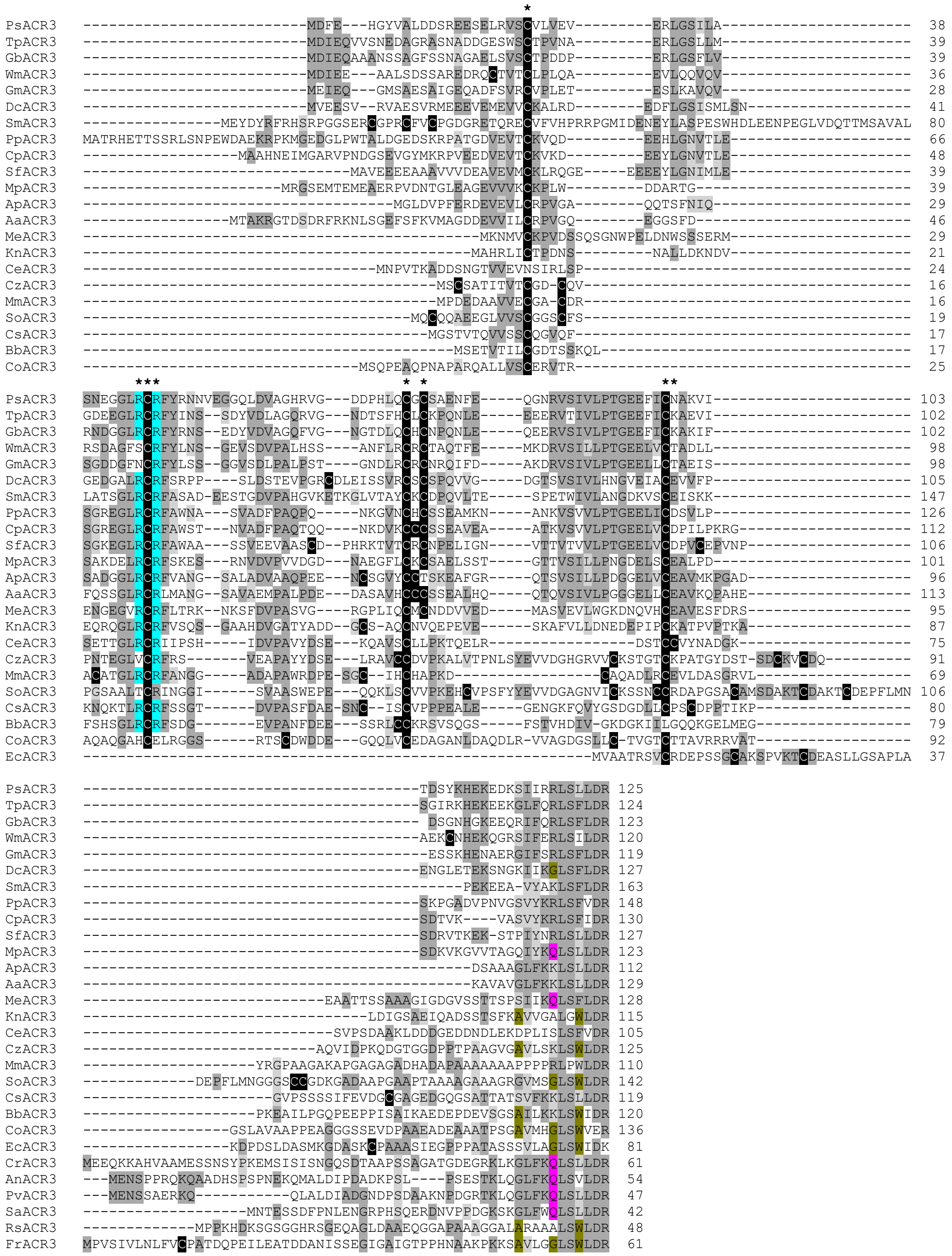

**Figure S3. Multiple sequence alignment of N-terminal regions of plant ACR3 proteins analyzed *in silico*.** At least 25% sequence identity in a column is highlighted. Cysteine residues are indicated in black. Di-arginine motifs are highlighted in blue. Conserved residues of MpACR3 analyzed in this work are marked with asterisks. Multiple sequence alignment was generated using the Clustal Omega server (https://www.ebi.ac.uk/Tools/msa/clustalo/) and manually adjusted. Sequence details are provided in Table S1.

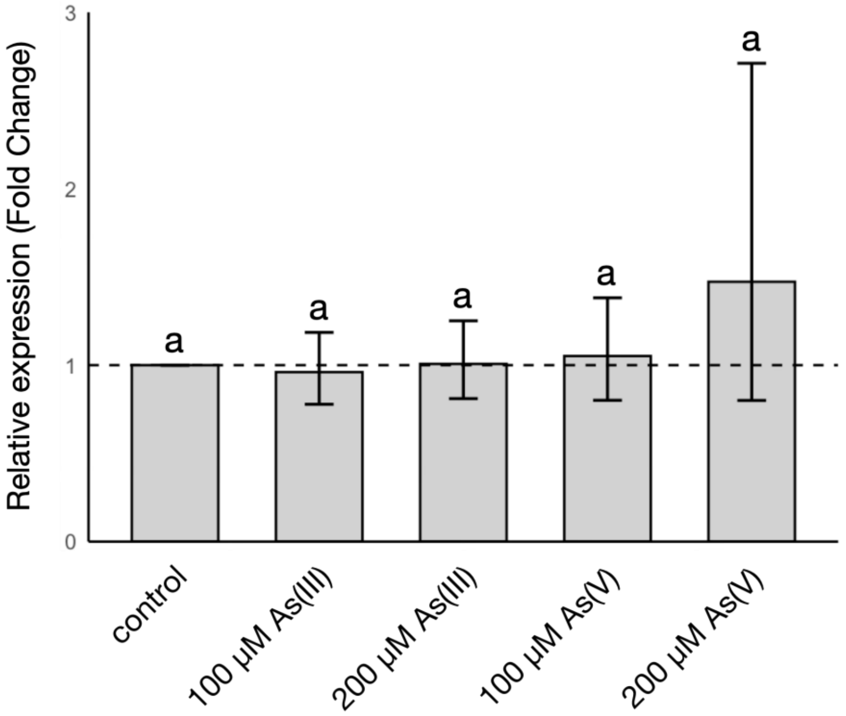

**Figure S4. Arsenic exposure does not affect *PpACR3* expression.** Total RNA was isolated from wild-type *Physcomitrium patens* shoots grown on solid BCD medium in the absence (control) or presence of 100 µM or 200 µM As(III) or As(V). *PpACT* (*ACTIN5*) was used as a reference gene for normalization of *PpACR3* expression levels. Data represent fold changes relative to the untreated control (set to 1.0 and indicated by a dashed horizontal line). Bars represent mean fold-change values from biological replicates for each treatment group (n = 6). Error bars indicate 95% confidence intervals. Values >1 indicate upregulation, whereas values <1 indicate downregulation relative to the control. Different letters indicate statistically significant differences as determined by the Kruskal–Wallis test followed by Dunn's multiple comparisons test ($P < 0.05$).

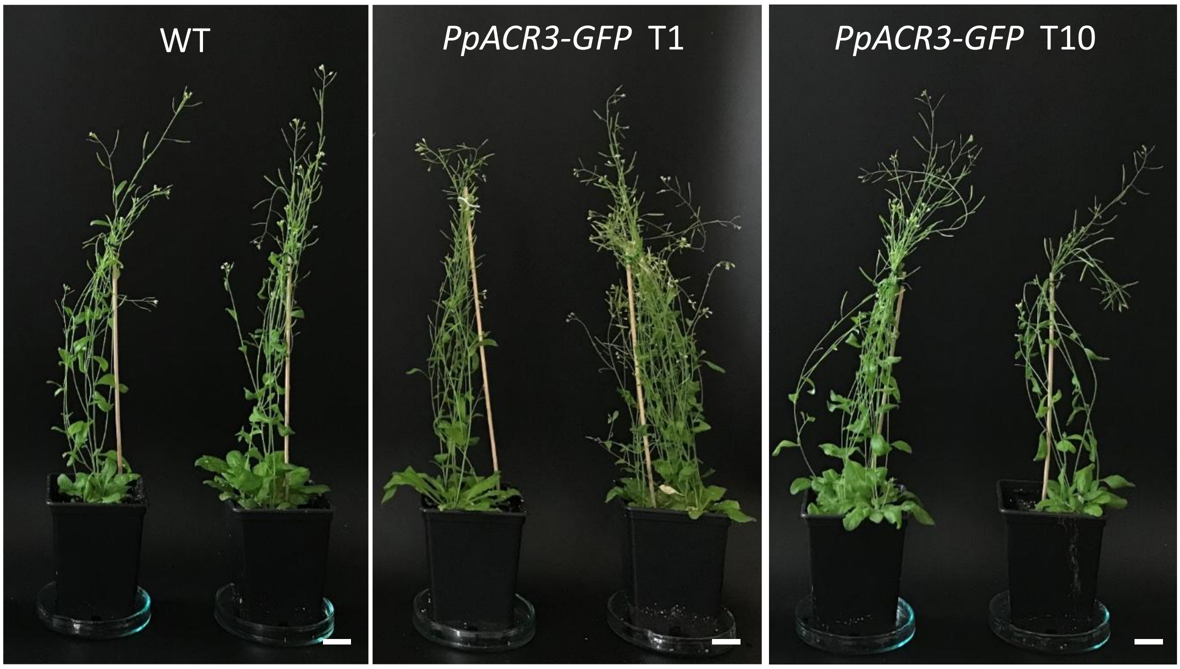

| *P. patens* lines | Main stem height (cm) | Number of side branches | Flowers - main stem | All flowers per stem | N |
|---|---|---|---|---|---|
| **WT** | 26 (+/-2.9) | 5.1 (+/- 1.3) | 20.8 (+/- 6.3) | 15.3 (+/- 6.3) | 8 |
| ***PpACR3-GFP* T10** | 22.7 (+/- 5.3) | 4 (+/- 1.1) | 19.3 (+/- 5.7) | 14.5 (+/-5.7) | 8 |
| ***PpACR3-GFP* T1** | 26.1 (+/- 4.9) | 4.6 (+/- 1.9) | 24 (+/- 6.6) | 16.4 (+/- 7.6) | 14 |

**Figure S5. No phenotypic differences were observed between wild-type (WT, Col-0) *Arabidopsis thaliana* plants and pro35S:PpACR3-GFP transgenic lines (T1 and T10) under control conditions.** Plants were grown in soil under long-day (LD; 16 h light/8 h dark, 22°C) conditions. The indicated growth parameters were measured or counted after completion of the growth cycle. Scale bars = 2 cm.

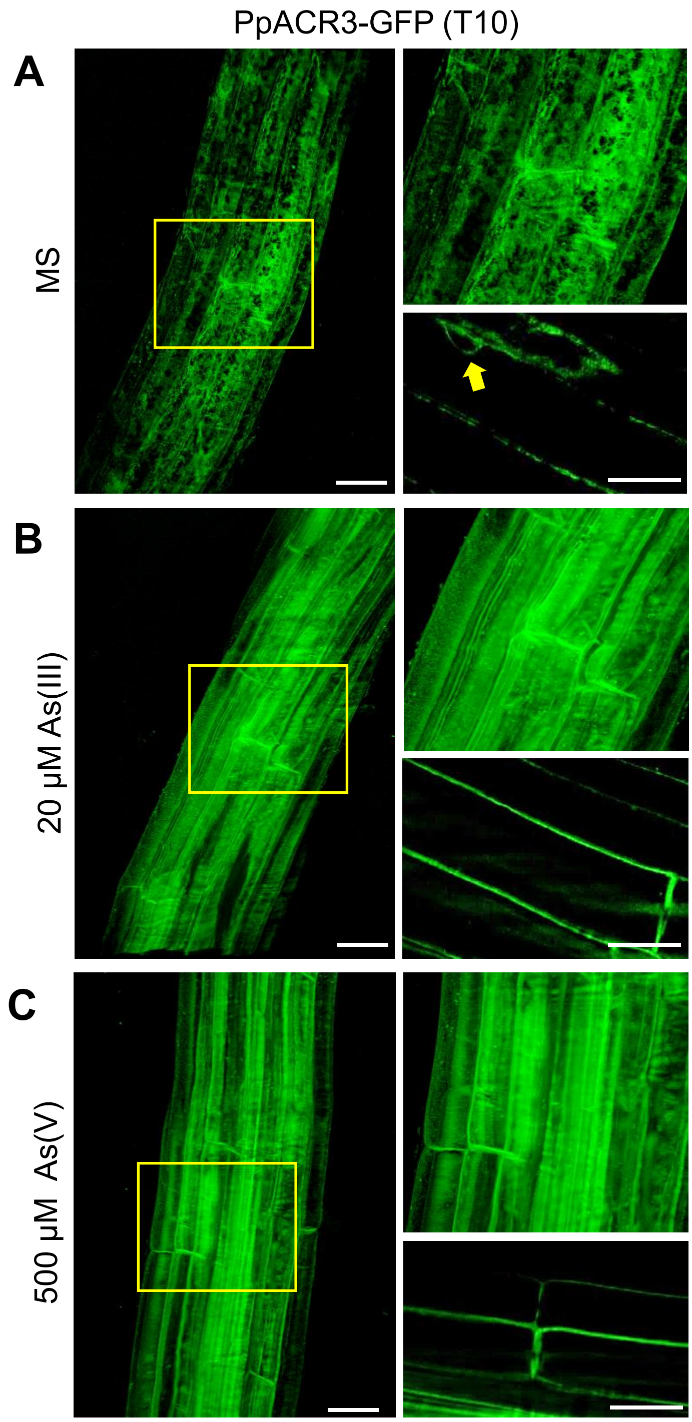

**Figure S6. Subcellular localization of PpACR3 in plant cells.** (A) PpACR3-GFP localized intracellularly in *A. thaliana* root cells (line T10) when grown under basal conditions (MS medium). Yellow arrow indicates the characteristic perinuclear signal. (B, C) In the presence of As(III) and As(V), PpACR3-GFP localized exclusively to the PM. PpACR3-GFP expressing seedlings (line T10) were grown for 4 days on MS medium and transferred to MS medium with or without As(III) or As(V) and grown for 24 h before imaging with the use of the ZEISS Lattice Lightsheet 7 microscopy. Roots are displayed in 3D space or as 2D views of single planes. Three plants from each treatment group were analyzed, and representative images are shown. Scale bars = 50 µm and 15 µm for 3D and 2D images, respectively.

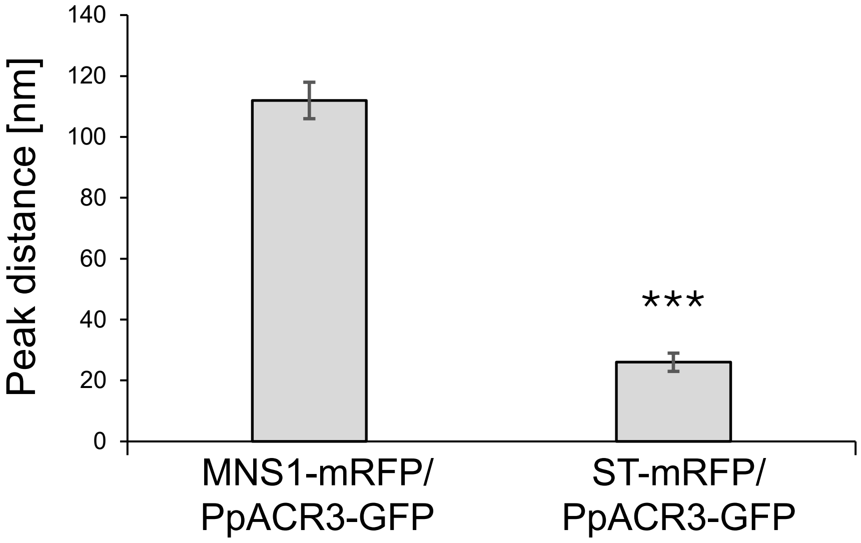

**Figure S7. Co-localization line profile analysis for PpACR3-GFP with *cis*- and *trans*-Golgi marker constructs.** Peak distance analysis indicates the distance between PpARC3-GFP and MNS1-mRFP = 112 ± 6 nm, whilst the distance between PpARC3-GFP and ST-mRFP = 26 ± 3 nm. Significance was analyzed by Kruskal–Wallis (***, $p < 0.001$). Results shown from n = 3 biological replicas with at least 6 technical repeats. For comparison the peak distance difference between MNS1-GFP with MNS1-mRFP is 18 ± 8 nm and MNS1-GFP with ST-mRFP is 122 ± 12 nm (McGinness et al. 2025).

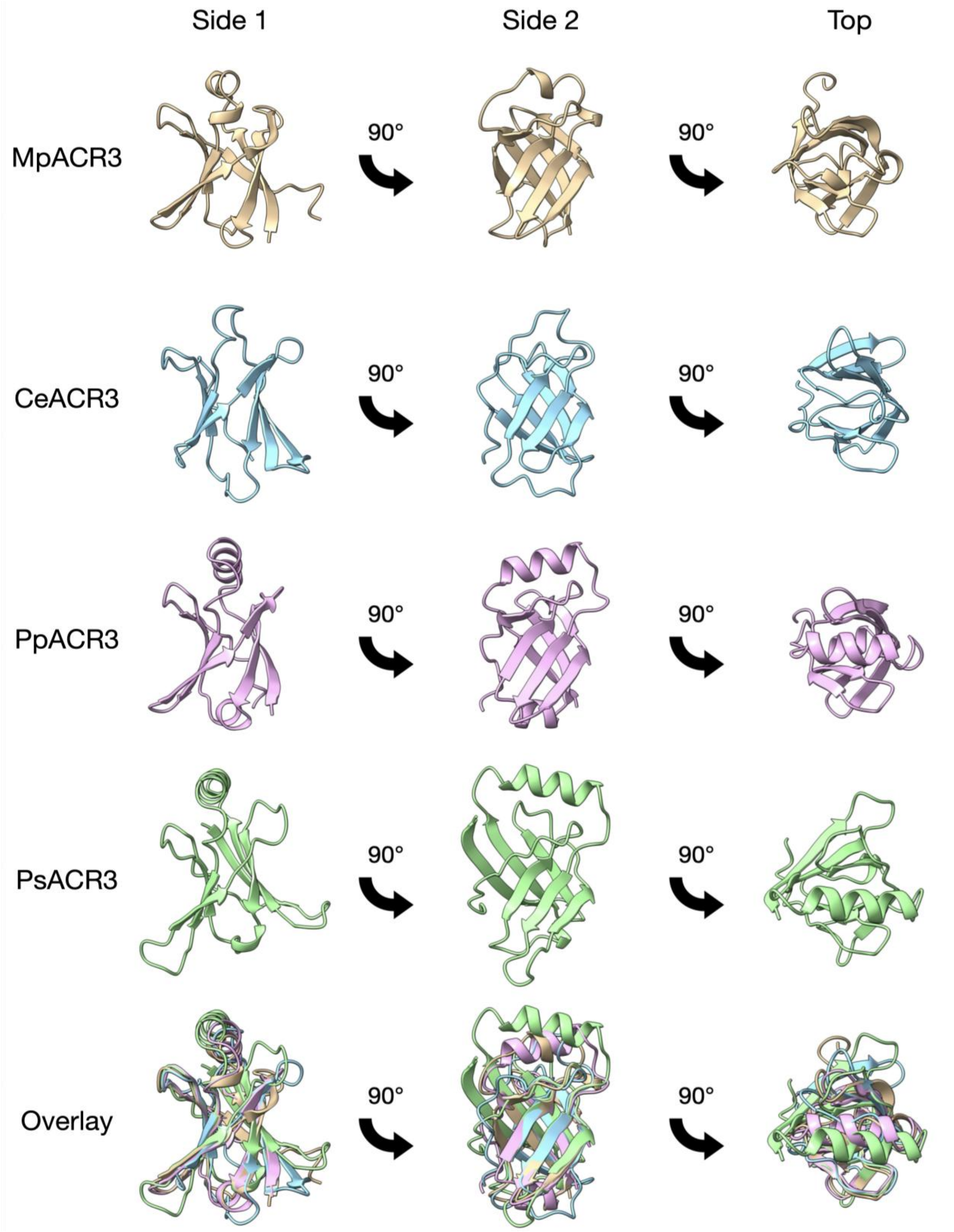

**Figure S8. Comparison of predicted N-terminal domain structures of selected plant ACR3 orthologs.** Structural models were generated using AlphaFold Server 3 (https://alphafoldserver.com) and visualized with UCSF ChimeraX 1.9 software (Meng et al., 2023)

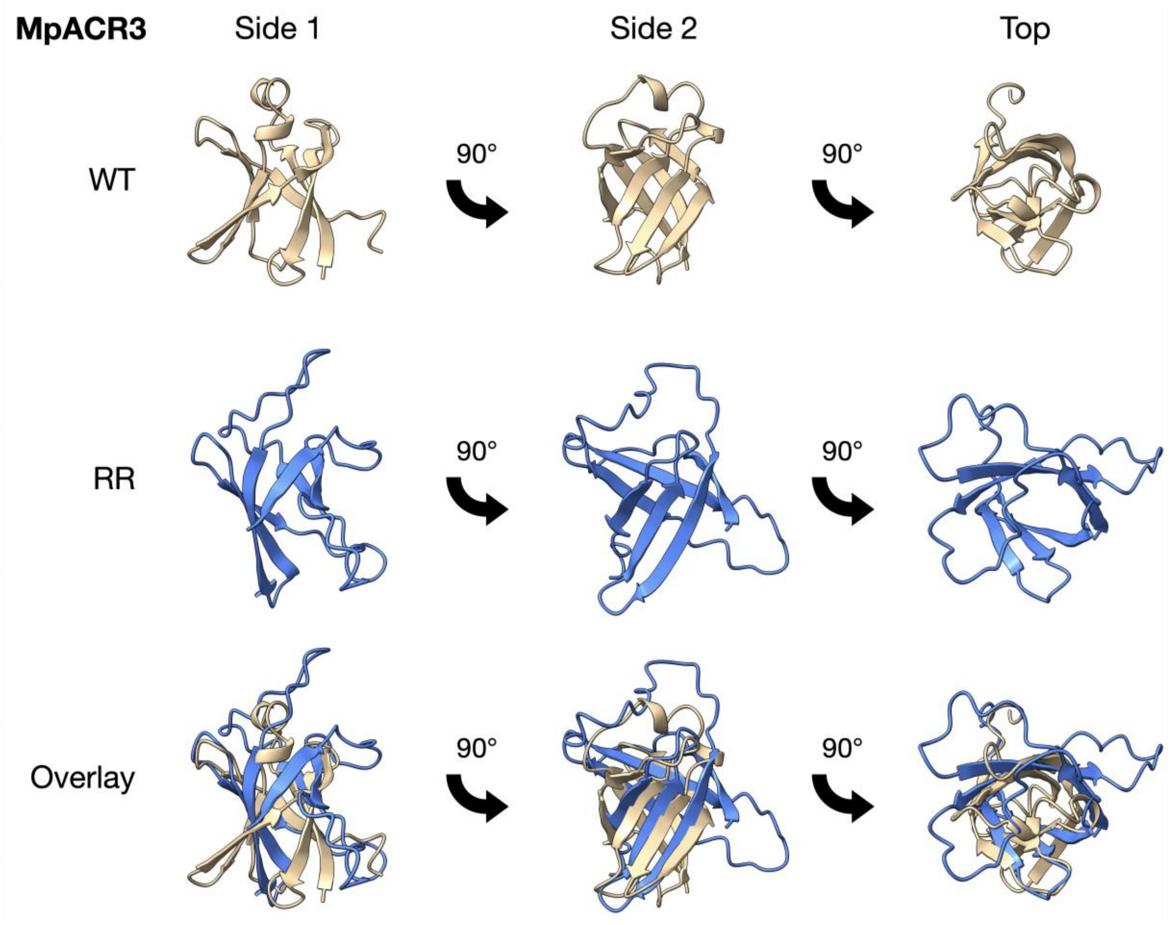

**Figure S9. Comparison of predicted N-terminal domain structures of the WT and RR variants of MpACR3.** Structural models were generated using AlphaFold Server 3 (https://alphafoldserver.com) and visualized with UCSF ChimeraX 1.9 (Meng et al., 2023).

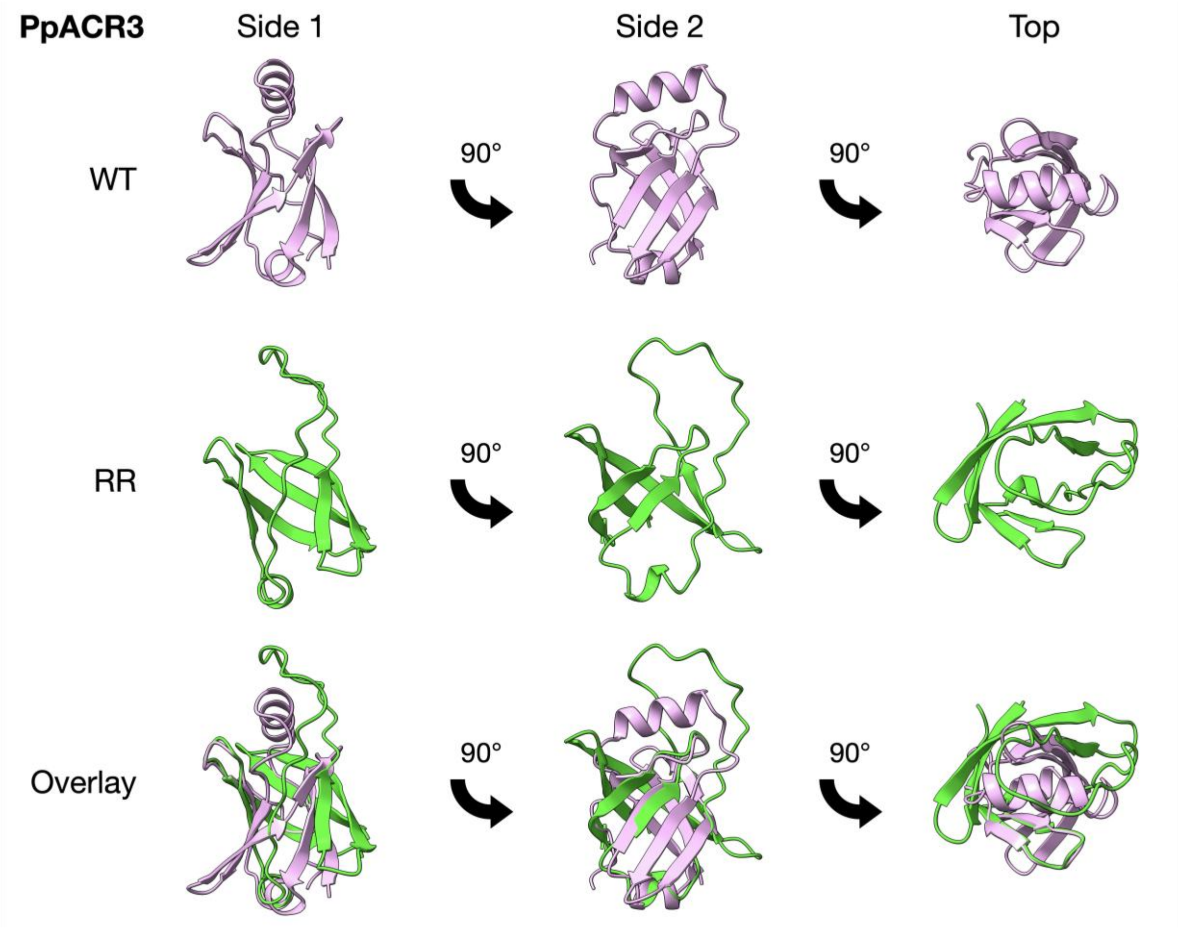

**Figure S10. Comparison of predicted N-terminal domain structures of the WT and RR variant of PpACR3.** Structural models were generated using AlphaFold Server 3 (https://alphafoldserver.com) and visualized with UCSF ChimeraX 1.9 (Meng et al., 2023).

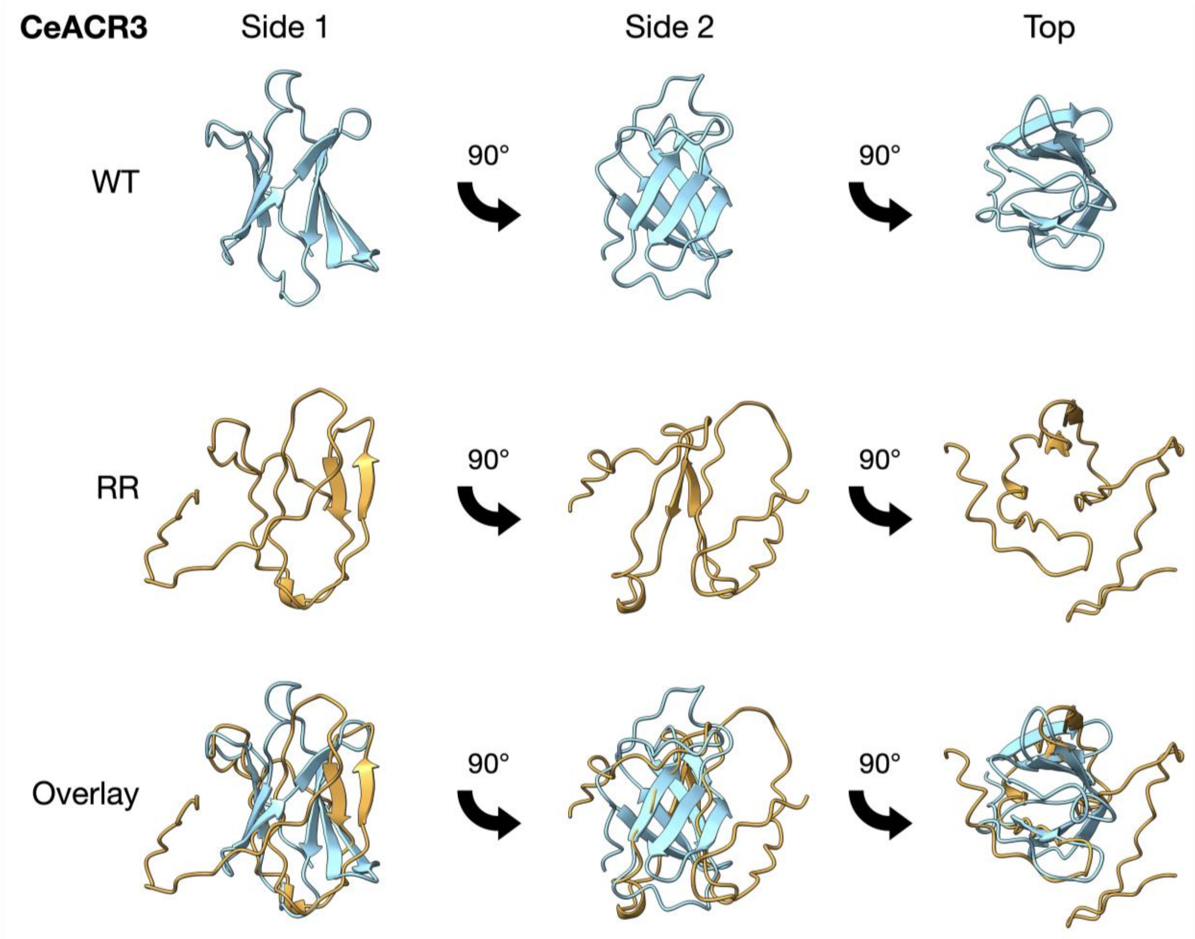

**Figure S11. Comparison of predicted N-terminal domain structures of the WT and RR variants of CeACR3.** Structural models were generated using AlphaFold Server 3 (https://alphafoldserver.com) and visualized with UCSF ChimeraX 1.9 (Meng et al., 2023).

**Table S1.** Plant ACR3 proteins analyzed *in silico* in this study.

| Protein name | Organism | Taxonomic group | Database | Protein identifier |
|---|---|---|---|---|
| PsACR3 | *Picea sitchensis* | Gymnosperms | NCBI[1] | ADE76924.1 |
| TpACR3 | *Thuja plicata* | Gymnosperms | Phytozome[2] | Thupl.29379155s0008 |
| GbACR3 | *Ginkgo biloba* | Gymnosperms | Gigascience[3] | Gb_32402 |
| WmACR3 | *Welwitschia mirabilis* | Gymnosperms | Data Dryad[4] | W.mirabilis.22940 |
| GmACR3 | *Gnetum montanum* | Gymnosperms | Data Dryad[5] | TnS000269885t05 |
| PvACR3 | *Pteris vittata* | Ferns | NCBI | ADP20955.1 |
| CrACR3 | *Ceratopteris richardii* | Ferns | NCBI | KAH7279060.1 |
| AnACR3 | *Adiantum nelumboides* | Ferns | NCBI | MCO5594610.1 |
| SaACR3 | *Salvinia cucullata* | Ferns | Fernbase[6] | Sacu_v1.1_s0024.g008994 |
| DcACR3 | *Diphasiastrum complanatum* | Clubmosses | NCBI | KAJ7529772.1 |
| SmACR3 | *Selaginella moellendorffii* | Spikemosses | NCBI | XP_024531516.1 |
| PpACR3 | *Physcomitrium patens* | Mosses | NCBI | XP_024362327.1 |
| CpACR3 | *Ceratodon purpureus* | Mosses | NCBI | KAG0559408.1 |
| SfACR3 | *Sphagnum fallax* | Mosses | NCBI | KAH8961229.1 |
| MpACR3 | *Marchantia polymorpha* | Liverworts | NCBI | OAE35577.1 |
| AaACR3 | *Anthoceros agrestis* | Hornworts | Hornwort genomes[7] | AagrOXF_evm.model.utg000091l.433__434.1 |
| ApACR3 | *Anthoceros punctatus* | Hornworts | Hornwort genomes | Apun_evm.model.utg000254l.51.1 |
| MeACR3 | *Mesotaenium endlicherianum* SAG 12.97 | Charophytes | Phycocosm[8] | https://phycocosm.jgi.doe.gov/cgi-bin/dispGeneModel?db=Mesen1_1&id=7825 |
| KnACR3 | *Klebsormidium nitens* NIES-2285 | Charophytes | NCBI | GAQ91548.1 |
| CsACR3 | *Coccomyxa subellipsoidea* C-169 | Chlorophytes | Phycocosm | https://phycocosm.jgi.doe.gov/cgi-bin/dispGeneModel?db=Cosub3&id=1208183 |
| CoACR3 | *Chlorella ohadii* isolate 1 | Chlorophytes | NCBI | KAI7841530.1 |
| BbACR3 | *Botryococcus braunii* Showa | Chlorophytes | Phycocosm | https://phycocosm.jgi.doe.gov/cgi-bin/dispGeneModel?db=Botrbrau1&id=23661 |
| CeACR3 | *Chlamydomonas eustigma* NIES-2499 | Chlorophytes | NCBI | GAX84956.1 |
| CzACR3 | *Chromochloris zofingiensis* | Chlorophytes | Phycocosm | https://phycocosm.jgi.doe.gov/cgi-bin/dispGeneModel?db=Chrzof1&id=14366 |
| MmACR3 | *Monoraphidium minutum* isolate 26B-AM | Chlorophytes | NCBI | KAI8476375.1 |
| RsACR3 | *Raphidocelis subcapitata* NIES-35 | Chlorophytes | NCBI | GBF99582.1 |
| SoACR3 | *Scenedesmus obliquus* UTEX 3031 | Chlorophytes | NCBI | WIA08162.1 |

| | | | | |
|---|---|---|---|---|
| EcACR3 | *Enallax costatus* CCAP 276/31 | Chlorophytes | Phycocosm | https://phycocosm.jgi.doe.gov/cgi-bin/dispGeneModel?db=Enacos1_1&id=6449963 |
| FrACR3 | *Flechtneria rotunda* SEV3VF49 | Chlorophytes | Phycocosm | https://phycocosm.jgi.doe.gov/cgi-in/dispGeneModel?db=Flerot1_1&id=11394391 |

[1]www.ncbi.nlm.nih.gov
[2]phytozome-next.jgi.doe.gov
[3]gigadb.org/dataset/100613
[4]datadryad.org/stash/dataset/doi:10.5061/dryad.ht76hdrdr
[5]datadryad.org/stash/dataset/doi:10.5061/dryad.0vm37
[6]fernbase.org
[7]www.hornworts.uzh.ch/en/hornwort-genomes.html
[8]phycocosm.jgi.doe.gov

**Table S2.** Characteristics of the plant species selected for this study.

| Species | Habitat type | Application | References |
| --- | --- | --- | --- |
| *C. subellipsoidea* | Terrestrial/freshwater (soil biofilms, subaerial habitats); free-living in diverse environments including cold/extreme conditions; tolerates variable hydration and temperature | Extremophile adaptation studies; biotechnology applications | Blanc et al., 2012<br>Kania et al., 2023 |
| *R. subcapitata* | Freshwater; ponds, lakes, rivers; temperate and tropical freshwater systems | Standard ecotoxicology test organism; model for environmental monitoring and chemical toxicity assessment | Suzuki et al., 2018 Yamagishi et al., 2020<br>Machado et al., 2024<br>Yeap et al., 2024 |
| *C. eustigma* | Freshwater; extremely acidic aquatic environments; high metal content; acid mine drainage sites | Model for acidophilic adaptation and heavy metal detoxification mechanisms | Hirooka et al., 2017 |
| *M. polymorpha* | Moist terrestrial; cosmopolitan distribution except Antarctica; pioneering species; moist soil, disturbed sites | Model plant for evolutionary studies, developmental biology and environmental stress research | Bowman et al., 2022<br>Poveda et al., 2025<br>Naramoto et al., 2022 |
| *P. patens* | Moist terrestrial; exposed mud, seasonally wet areas; temperate regions worldwide | Model plant for evolutionary studies and developmental biology | Rensig et al., 2020 Naramoto et al., 2022 |
| *P. vittata* | Terrestrial; arsenic-contaminated soils; disturbed sites; tropical and subtropical regions | Arsenic hyperaccumulator; phytoremediation applications | Danh et al., 2014<br>Bai et al., 2023 |
| *P. sitchensis* | Coastal temperate forests | Forestry and climate adaptation research | Elleouet et al., 2019 Gagalova et al., 2022 |

**Table S3.** Pairwise sequence identity and similarity among plant ACR3 orthologs.

| Identity | | | | | | | | Identity stats | |
|---|---|---|---|---|---|---|---|---|---|
| PvACR3 | 100% | | | | | | | min | 41.5 |
| CsACR3 | 56.11% | 100% | | | | | | max | 65.2 |
| RsACR3 | 53.29% | 61.38% | 100% | | | | | mean | 54.3 |
| MpACR3 | 65.17% | 57.22% | 52.88% | 100% | | | | stddev | 6.2 |
| CeACR3 | 50.65% | 59.72% | 50.62% | 43.76% | 100% | | | | |
| PpACR3 | 57.78% | 54.72% | 49.37% | 56.45% | 41.50% | 100% | | | |
| PsACR3 | 64.90% | 56.11% | 50.62% | 59.42% | 44.51% | 54.38% | 100% | | |
| | PvACR3 | CsACR3 | RsACR3 | MpACR3 | CeACR3 | PpACR3 | PsACR3 | | |
| **Similarity** | | | | | | | | **Similarity stats** | |
| PvACR3 | 100% | | | | | | | min | 51.6 |
| CsACR3 | 69.16% | 100% | | | | | | max | 74.9 |
| RsACR3 | 63.58% | 71.94% | 100% | | | | | mean | 70.3 |
| MpACR3 | 74.93% | 68.05% | 62.90% | 100% | | | | stddev | 13.3 |
| CeACR3 | 62% | 69.44% | 62.65% | 54.48% | 100% | | | | |
| PpACR3 | 70.71% | 67.22% | 61.40% | 68.49% | 51.57% | 100% | | | |
| PsACR3 | 74.14% | 68.88% | 61.65% | 67.98% | 56.35% | 65.78% | 100% | | |
| | PvACR3 | CsACR3 | RsACR3 | MpACR3 | CeACR3 | PpACR3 | PsACR3 | | |

stddev, standard deviation;

**Table S4.** Predicted topology of plant ACR3 transporters.

| Protein name | Protein length | Evidence | Reliability | Number of TMs | Localization of N-terminus | Localization of C-terminus |
|---|---|---|---|---|---|---|
| CsACR3 | 360 | Prediction | 85 | 10 | Intracellular | Intracellular |
| RsACR3 | 399 | Prediction | 86 | 10 | Intracellular | Intracellular |
| PvACR3 | 379 | Prediction | 90 | 10 | Intracellular | Intracellular |
| MpACR3 | 457 | Prediction | 91 | 10 | Intracellular | Intracellular |
| CeACR3 | 495 | Prediction | 78 | 10 | Intracellular | Intracellular |
| PpACR3 | 477 | Prediction | 83 | 10 | Intracellular | Intracellular |
| PsACR3 | 456 | Prediction | 92 | 10 | Intracellular | Intracellular |

TM, transmembrane region;

**Table S5.** Oligonucleotides used in this work.

| Primer name | 5' to 3' sequence |
|---|---|
| **Gene cloning:** | |
| BamHI-CsACR3-fw | TAGAACTAGTGGATCCATGAGATCTGCTGGTGTTTTTAAG |
| SalI-CsACR3-rv | GACATGTCGAGGTCGACAGCAGAAGCACCAGTAGTACCT |
| BamHI-RsACR3-fw | ATACTCTAGAACTAGTGGATCCATGCCACCAAAACACGATAAGTCTG |
| SalI-RsACR3-rv | CTTTAGACATGTCGAGGTCGACCTTAACAGCACCAGCTGTAGAACC |
| BamHI-CeACR3-fw | ATACTCTAGAACTAGTGGATCCATGAATCCAGTTACTAAGG |
| SalI-CeACR3-rv | CTTTAGACATGTCGAGGTCGACTGGCTTGATCAGTTGAGCAACAG |
| BamHI-MpACR3-fw | TAGAACTAGTGGATCCATGAGGGGCTCGGAGATG |
| SalI-MpACR3-rv | ACATGTCGAGGTCGACAGCTTGTTCTTTTGAGAGCC |
| BamHI-PpACR3-fw | TAGAACTAGTGGATCCATGGCTACTAGACACGAAACTACA |
| SalI-PpACR3-rv | ACATGTCGAGGTCGACGAAAAACTTCTTACGGAAGAACAAA |
| BamHI-PsACR3-fw | TAGAACTAGTGGATCCATGGACTTTGAACATGGTTAGTTG |
| SalI-PsACR3-rv | ACATGTCGAGGTCGACACCGAACCACTTAGTTTTCAAGGC |
| PpACR3-GUSGFP-fw | GGAGAGAACACGGGGGACTCTAGAATGGCAACGCGTCACGAGACAACG |
| PpACR3-GUSGFP-rv1 | TGGGATCGAATTGATCCTCTAGGAAGAATTTCTTCCTAAAAAAGAGAGAAACG |
| **Sequencing:** | |
| MET17_fw | GCGTCTGTTAGAAAGGAAGT |
| yeGFP_rv | GTGACCATTAACATCACCATC |
| **Mutagenesis:** | |
| CeC32A-fw | GAAACTACCGGTTTGAGAGCCAGAATTATCCCATCTCATATTG |
| CeC32A-rv | CAATATGAGATGGGATAATTCTGGCTCTCAAACCGGTAGTTTC |
| CeC54A-fw | CTGAAAAGCAAGCTGTTTCTGCCTTGTTGCCAAAGACTCAAG |
| CeC54A-rv | CTGAAAAGCAAGCTGTTTCTGCCTTGTTGCCAAAGACTCAAG |
| CeC67A-fw | CAAGAATTGAGAGATTCTACCGCTTGCGTTTACAACGCTGATGG |
| CeC67A-rv | CCATCAGCGTTGTAAACGCAAGCGGTAGAATCTCTCAATTCTTG |
| CeC68A-fw | TCAAGAATTGAGAGATTCTACCTGTGCCGTTTACAACGCTGATGG |
| CeC68A-rv | CCATCAGCGTTGTAAACGGCACAGGTAGAATCTCTCAATTCTTGA |
| CeRR-fw | GAAACTACCGGTTTGGCATGCGCAATTATCCCATCTCATATTG |
| CeRR-rv | CAATATGAGATGGGATAATTGCGCATGCCAAACCGGTAGTTTC |
| CeRCR-fw | TCCATCTGAAACTACCGGTTTGGCAGCCGCAATTATCCCAT |
| CeRCR-rv | ATGGGATAATTGCGGCTGCCAAACCGGTAGTTTCAGATGGA |
| PpC52A-fw | TACTGGTAACGTTGAAGTTACCGCTAAAGTTCAAGACGAAGAACAC |
| PpC52A-rv | GTGTTCTTCGTCTTGAACTTTAGCGGTAACTTCAACGTTACCAGTA |
| PpC74A-fw | TCTGGTAGAGAAGGTTTGAGAGCTAGATTTGCTTGGAATGCTTC |
| PpC74A-rv | GAAGCATTCCAAGCAAATCTAGCTCTCAAACCTTCTCTACCAGA |
| PpC96A-fw | GCTCAACCACAAAACAAGGGTGTTAATGCTCATTGCTCATCCG |
| PpC96A-rv | CGGATGAGCAATGAGCATTAACACCCTTGTTTTGTGGTTGAGC |
| PpC98A-fw | ACCACAAAACAAGGGTGTTAATTGTCATGCCTCATCCGAAGCTAT |
| PpC98A-rv | ATAGCTTCGGATGAGGCATGACAATTAACACCCTTGTTTTGTGGT |
| PpC121A-fw | GTTTTGCCAACCGGTGAAGAATTGATTGCCGATTCTGTTTTGCC |
| PpC121A-rv | GGCAAAACAGAATCGGCAATCAATTCTTCACCGGTTGGCAAAAC |
| PpRR-fw | TTGGAATCTGGTAGAGAAGGTTTGGCATGTGCATTTGCTTGGAATGCTTCTGTTGC |
| PpRR-rv | GCAACAGAAGCATTCCAAGCAAATGCACATGCCAAACCTTCTCTACCAGATTCCAA |
| MpRR-fw | AGCGCGAAGGATGAGTTGGCGTGTGCGTTCTCCAAGGAATCGAG |

| | |
|---|---|
| MpRR-rv | CTCGATTCCTTGGAGAACGCACACGCCAACTCATCCTTCGCGCT |
| **qPCR** | |
| PpACR3_RT-fw2 | CAAGATGAGGAGACTTGGGG |
| PpACR3_RT-rv2 | ACACCCTTGTTCTGGGGTTG |
| PpACT-fw | ACCGAGTCCAACATTCTACC |
| PpACT-rv | GTCCACATTAGATTCTCGCA |

**Table S6.** Plasmids used in this work.

| Name | Description | Source |
|---|---|---|
| pUG35 | MET17 promoter, yeGFP, CEN, URA3, AmpR | Lab collection |
| pScACR3 | $_{pro}$*MET17:ScACR3-yeGFP* in pUG35 | Mizio et al. (2023) |
| pCsACR3 | $_{pro}$*MET17:CsACR3-yeGFP* in pUG35 | This study |
| pRsACR3 | $_{pro}$*MET17:RsACR3-yeGFP* in pUG35 | This study |
| pCeACR3 | $_{pro}$*MET17:CeACR3-yeGFP* in pUG35 | This study |
| pMpACR3 | $_{pro}$*MET17:MpACR3-yeGFP* in pUG35 | Mizio et al. (2025) |
| pPpACR3 | $_{pro}$*MET17:PpACR3-yeGFP* in pUG35 | This study |
| pPvACR3 | $_{pro}$*MET17:PvACR3-yeGFP* in pUG35 | This study |
| pPsACR3 | $_{pro}$*MET17:PsACR3-yeGFP* in pUG35 | This study |
| pCeACR3-C32A | *proMET17:CsACR3-C32A-yeGFP* in pUG35 | This study |
| pCeACR3-C54A | *proMET17:CsACR3-C54A-yeGFP* in pUG35 | This study |
| pCeACR3-C67A | *proMET17:CsACR3-C67A-yeGFP* in pUG35 | This study |
| pCeACR3-C68A | *proMET17:CsACR3-C68A-yeGFP* in pUG35 | This study |
| pCeACR3-RR | *proMET17:CsACR3-R31A,R33A-yeGFP* in pUG35 | This study |
| pCeACR3-RCR | *proMET17:CsACR3-R31A,C32A,R33A-yeGFP* in pUG35 | This study |
| pPpACR3-C52A | *proMET17:PpACR3-C52A-yeGFP* in pUG35 | This study |
| pPpACR3-C74A | *proMET17:PpACR3-C74A-yeGFP* in pUG35 | This study |
| pPpACR3-C96A | *proMET17:PpACR3-C96A-yeGFP* in pUG35 | This study |
| pPpACR3-C98A | *proMET17:PpACR3-C98A-yeGFP* in pUG35 | This study |
| pPpACR3-C121A | *proMET17:PpACR3-C121A-yeGFP* in pUG35 | This study |
| pPpACR3-RR | *proMET17:PpACR3-R73A,R75A-yeGFP* in pUG35 | This study |
| pMpACR3-RR | *proMET17:MpACR3-R46A,R48A-yeGFP* in pUG35 | This study |
| pGFPGUSPlus | CaMV 35S promoter, EGFP, GUSplus$^{TM}$, HygR, KanR | Vickers et al., 2007 |
| pGFPGUSPlus-PpACR3 | $_{pro}$*35S:PpACR3-EGFP* in pGFPGUSPlus | This study |

**Table S7.** Model scores and seed numbers for the N-terminal domains of the indicated ACR3 proteins.

| | MpACR3 | | CeACR3 | | PpACR3 | | PsACR3 | |
|---|---|---|---|---|---|---|---|---|
| **Seed** | **pTM** | **pLDDT** | **pTM** | **pLDDT** | **pTM** | **pLDDT** | **pTM** | **pLDDT** |
| 2137 | 0.74 | | 0.7 | 75.98 | 0.71 | 76.10 | 0.73 | |
| 107231682 | 0.73 | | 0.69 | | 0.7 | | 0.73 | |
| 249895374 | **0.75** | **78.80** | 0.7 | 75.39 | 0.71 | 76.18 | 0.74 | 79.01 |
| 322293752 | 0.74 | | 0.7 | 75.05 | 0.7 | | 0.74 | 79.06 |
| 397615034 | 0.73 | | 0.7 | 75.65 | 0.7 | | 0.73 | |
| 730029121 | 0.74 | | 0.69 | | 0.69 | | 0.73 | |
| 775995308 | 0.74 | | 0.7 | 76.01 | 0.7 | | 0.74 | 78.50 |
| 862731019 | 0.74 | | 0.7 | 75.50 | 0.7 | | 0.74 | 77.62 |
| 1005710457 | 0.75 | 78.68 | 0.7 | 75.57 | 0.69 | | 0.74 | 78.39 |
| 1157336657 | 0.74 | | 0.7 | 75.59 | 0.7 | | 0.74 | 78.04 |
| 1349946916 | 0.75 | 78.72 | 0.7 | 75.83 | 0.7 | | 0.74 | 79.86 |
| 1414339364 | 0.74 | | 0.7 | 75.80 | 0.7 | | 0.74 | 78.53 |
| 1598553664 | 0.75 | 78.07 | 0.7 | 75.71 | 0.7 | | 0.74 | 76.57 |
| 1720798967 | 0.73 | | 0.7 | 75.19 | **0.71** | **76.49** | **0.74** | **80.44** |
| 1977933808 | 0.74 | | 0.7 | 76.02 | 0.7 | | 0.74 | 79.18 |
| 2046719219 | 0.75 | 78.12 | 0.69 | | 0.7 | | 0.74 | 76.95 |
| 2067571921 | 0.75 | 77.81 | 0.7 | 75.70 | 0.7 | | 0.72 | |
| 2147478647 | 0.74 | | **0.7** | **76.10** | 0.7 | | 0.74 | 79.44 |
| 80213153 | 0.74 | | 0.7 | 75.22 | 0.71 | 74.37 | 0.73 | |
| 559621 | 0.75 | 78.59 | 0.7 | 75.51 | 0.7 | | 0.73 | |

pTM, predicted Template Modeling score; pLDDT, predicted Local Distance Difference Test score. Models with the highest pTM values and subsequently selected based on pLDDT values are highlighted in yellow. Final models used in this study are highlighted in green.
